# Glutamine deprivation triggers NAGK-dependent hexosamine salvage

**DOI:** 10.1101/2020.09.13.294116

**Authors:** Sydney L. Campbell, Clementina Mesaros, Luke Izzo, Hayley Affronti, Michael Noji, Bethany E. Schaffer, Tiffany Tsang, Kathryn Sun, Sophie Trefely, Salisa Kruijning, John Blenis, Ian A. Blair, Kathryn E. Wellen

**Affiliations:** Department of Cancer Biology, University of Pennsylvania, Philadelphia, PA 19104; Abramson Family Cancer Research Institute, University of Pennsylvania, Philadelphia, PA 19104; Department of Systems Pharmacology and Translational Therapeutics, University of Pennsylvania, Philadelphia, PA 19104; Meyer Cancer Center and Department of Pharmacology, Weill Cornell Medicine, New York, NY 10021; Pancreatic Cancer Research Center, Perelman School of Medicine, University of Pennsylvania, Philadelphia, PA 19104; Center for Metabolic Disease Research, Department of Microbiology and Immunology, Lewis Katz School of Medicine, Temple University, Philadelphia, PA 19140

## Abstract

Tumors frequently exhibit aberrant glycosylation, which can impact cancer progression and therapeutic responses. The hexosamine biosynthesis pathway (HBP) produces uridine diphosphate N-acetylglucosamine (UDP-GlcNAc), a major substrate for glycosylation in the cell. Prior studies have identified the HBP as a promising therapeutic target in pancreatic ductal adenocarcinoma (PDA). The HBP requires both glucose and glutamine for its initiation. The PDA tumor microenvironment is nutrient poor, however, prompting us to investigate how nutrient limitation impacts hexosamine synthesis. Here, we identify that glutamine limitation in PDA cells suppresses de novo hexosamine synthesis but results in increased free GlcNAc abundance. GlcNAc salvage via N-acetylglucosamine kinase (NAGK) is engaged to feed UDP-GlcNAc pools. *NAGK* expression is elevated in human PDA, and *NAGK* deletion from PDA cells impairs tumor growth in mice. Together, these data identify an important role for NAGK-dependent hexosamine salvage in supporting PDA tumor growth.

## Introduction

Altered glycosylation is frequently observed in malignancies, impacting tumor growth as well as immune and therapeutic responses (Akella et al., 2019; Mereiter et al., 2019; Munkley, 2019). Several types of glycosylation, including O-GlcNAcylation and N-linked glycosylation, are dependent on the glycosyl donor uridine diphosphate N-acetylglucosamine (UDP-GlcNAc), which is synthesized by the hexosamine biosynthesis pathway (HBP). The HBP branches off from glycolysis with the transfer of glutamine’s amido group to fructose-6-phosphate (F-6-P) to generate glucosamine-6-phosphate (GlcN-6-P), mediated by the rate limiting enzyme glutamine—fructose-6-phosphate transaminase (GFPT1/2). The pathway further requires acetyl-CoA, ATP, and uridine triphosphate (UTP) to ultimately generate UDP-GlcNAc. O-GlcNAcylation, the addition of a single N-acetylglucosamine (GlcNAc) moiety onto a serine or threonine residue of intracellular proteins, is upregulated in multiple cancers (Akella, et al., 2019). Targeting O-GlcNAcylation suppresses the growth of breast, prostate, and colon cancer tumors (Caldwell et al., 2010; Ferrer et al., 2017; Gu et al., 2010; Guo et al., 2017; Lynch et al., 2012). Similarly, highly branched N-glycan structures are sensitive to HBP flux and are upregulated in malignant tissue, and targeting the relevant Golgi GlcNAc transferase enzymes can limit tumor growth and metastasis in vivo (Granovsky et al., 2000; Li et al., 2008; Zhou et al., 2011). Thus, improved understanding the regulation of the HBP in cancer could point towards novel therapeutic strategies.

Pancreatic ductal adenocarcinoma (PDA) is a deadly disease with a 5-year survival rate of 9% and a rising number of annual deaths (Rahib et al., 2014) (ACS Cancer Facts and Figures 2019, NIH SEER report 2019). Mutations in *KRAS* occur in nearly all cases of human PDA and drive extensive metabolic reprogramming in cancer cells. Enhanced flux into the HBP was identified as a primary metabolic feature mediated by mutant KRAS in PDA cells (Ying et al., 2012). Hypoxia, a salient characteristic of the tumor microenvironment (Lyssiotis and Kimmelman, 2017), was shown to further promote expression of glycolysis and HBP genes in pancreatic cancer cells (Guillaumond et al., 2013). Notably, the glutamine analog 6-diazo-5-oxo-L-norleucine (DON), which inhibits the HBP, suppressed PDA metastasis and sensitized PDA tumors to anti-PD1 therapy (Sharma et al., 2020). DON has also been reported to sensitize PDA cells to the chemotherapeutic gemcitabine in vitro (Chen et al., 2017). Additionally, a recently developed inhibitor targeting the HBP enzyme phosphoacetylglucosamine mutase 3 (PGM3) enhances gemcitabine-mediated reduction of xenograft tumor growth in vivo (Ricciardiello et al., 2020). Thus, the HBP may represent a therapeutic target in PDA, although the regulation of UDP-GlcNAc synthesis and the optimal strategies to target this pathway for therapeutic benefit in PDA remain poorly understood.

An outstanding question is the impact of the tumor microenvironment on UDP-GlcNAc synthesis. The HBP has been proposed as a nutrient-sensing pathway since its rate-limiting step, mediated by GFPT1/2, requires both glutamine and the glycolytic intermediate fructose-6-phosphate (Denzel and Antebi, 2015). In hematopoietic cells, glucose deprivation limits UDP-GlcNAc levels and dramatically reduces levels of the N-glycoprotein IL3Rα at the plasma membrane in a manner dependent on the HBP (Wellen et al., 2010). Similarly, O-GlcNAcylation of certain nuclear-cytosolic proteins, including cancer-relevant proteins such as Myc and Snail, has been demonstrated to be nutrient sensitive, impacting protein stability or function (Housley et al., 2008; Park et al., 2010; Swamy et al., 2016). Yet, the PDA tumor microenvironment is thought to be particularly nutrient poor, owing to its characteristic dense stroma (Halbrook and Lyssiotis, 2017). This raises the question of how nutrient deprivation impacts the synthesis of UDP-GlcNAc and its utilization for glycosylation. Understanding how PDA cells regulate these processes under nutrient limitation could identify therapeutic vulnerabilities. In this study, we investigated the impact of nutrient deprivation on the HBP and glycosylation in PDA cells, identifying a key role for hexosamine salvage through the enzyme N-acetylglucosamine kinase (NAGK) in PDA tumor growth.

## Materials and methods

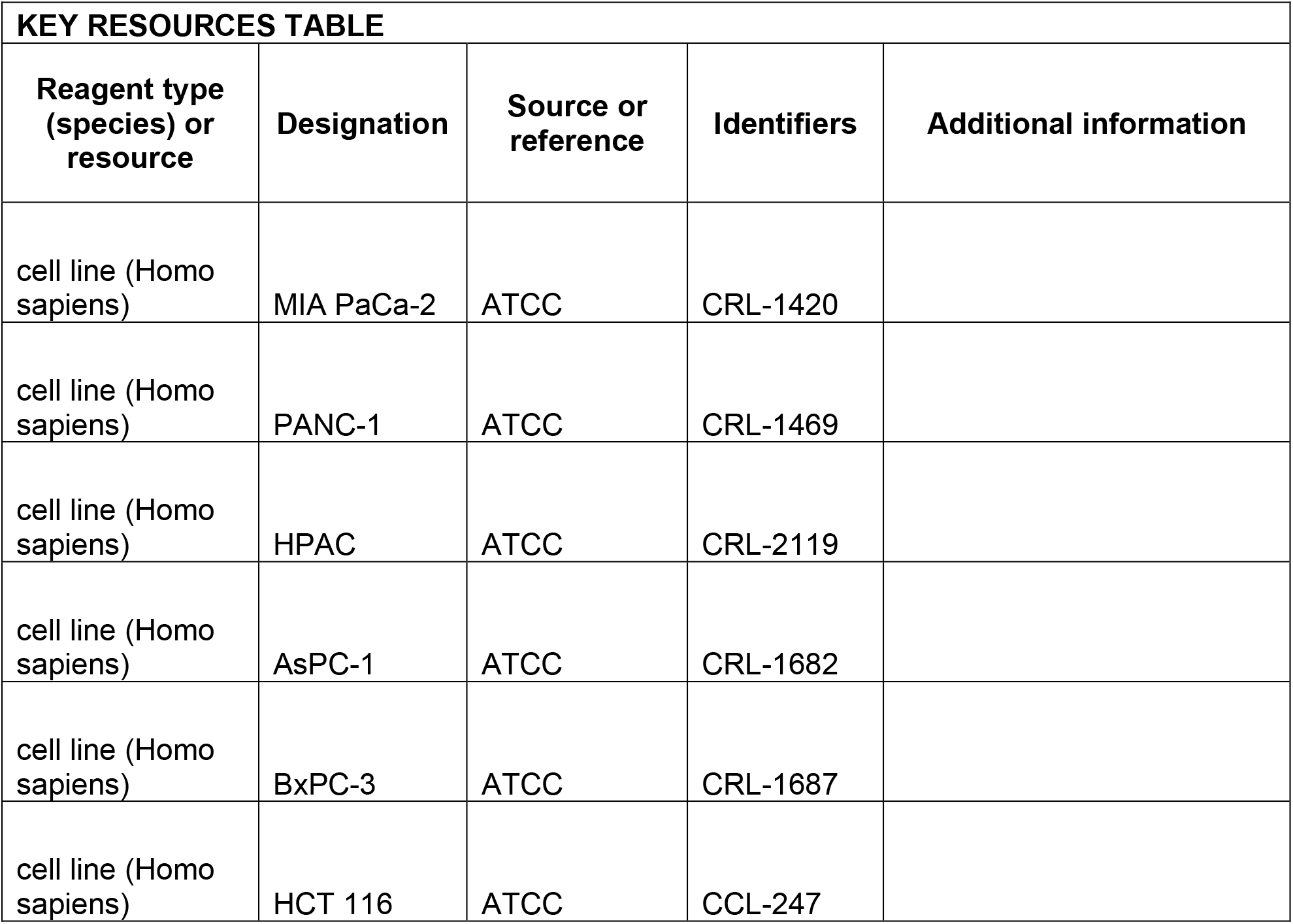
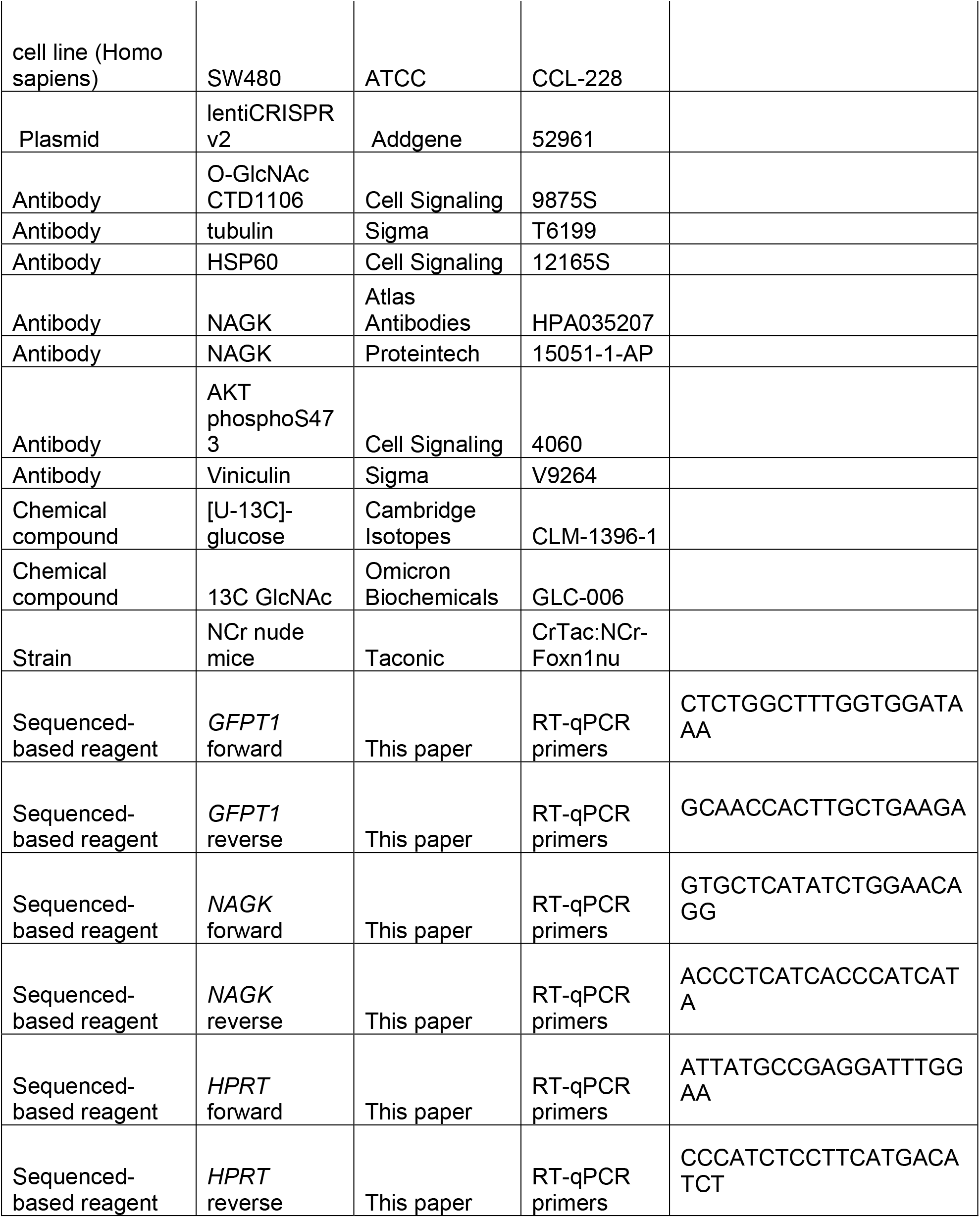
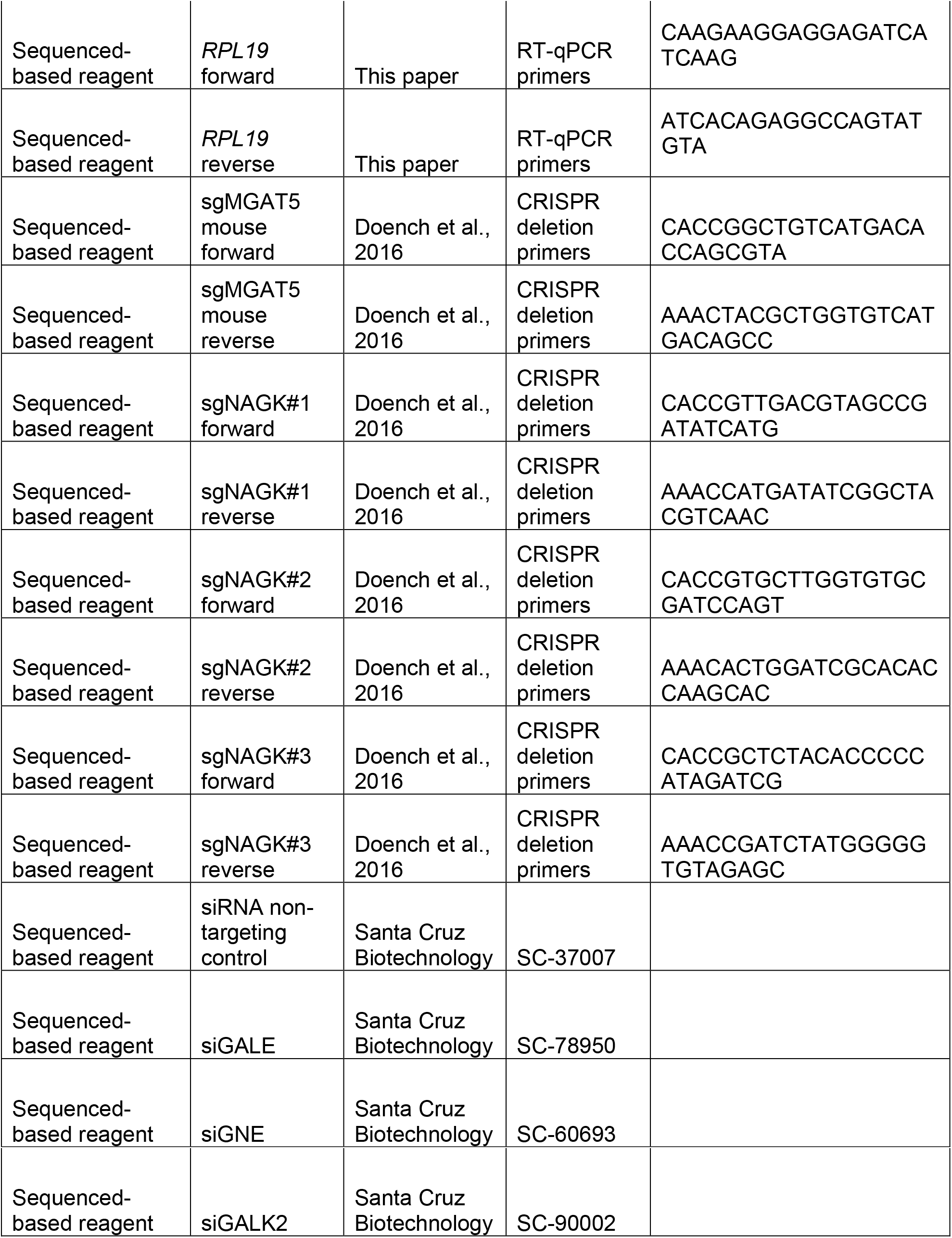

### Cell culture

Cells were cultured in DMEM high glucose (Gibco, 11965084) with 10% calf serum (Gemini GemCell U.S. Origin Super Calf Serum, 100-510), unless otherwise noted. Glucose- or glutamine-restricted media was prepared using glucose, glutamine, and phenol red free DMEM (Gibco, A1443001) supplemented with glucose (Sigma-Aldrich, G8769), glutamine (Gibco, 25030081), and dialyzed fetal bovine serum (Gemini, 100-108). For all glutamine restriction experiments except S2.3 D, cells were plated 2-3x more densely for the nutrient restricted condition samples to achieve similar confluency at the experiment endpoint. 1% oxygen levels were achieved by culturing cells in a Whitley H35 Hypoxystation (Don Whitley Scientific). ATCC names and numbers for the cell lines used in this study are: MIA PaCa-2 (ATCC# CRL-1420), PANC-1 (ATCC# CRL-1469), HPAC (ATCC# CRL-2119), AsPC-1 (ATCC# CRL-1682), BxPC-3 (ATCC# CRL-1687), HCT 116 (ATCC# CCL-247), and SW480 (ATCC# CCL-228). All cells were routinely tested for mycoplasma and authenticated by short tandem repeat (STR) profiling using the GenePrint 10 System (Promega, B9510).

### Generation of CRISPR cell lines

sgRNA sequences targeting *NAGK* or *Mgat5* from the Brunello and Brie libraries (Doench et al., 2016) were cloned into the lentiCRISPRv2 vector (Sanjana, Shalem et al., 2014). Lentivirus was produced in 293T cells according to standard protocol. Cells were then infected with the CRISPR lentivirus and selected with puromycin. Cells were plated at very low density into 96 well plates to establish colonies generated from single cell clones. *Mgat5* gene disruption was validated by qPCR and L-PHA binding. *NAGK* gene disruption was validated by qPCR, western blot, and ^13^C-GlcNAc tracing. Seven *NAGK* knockout clonal cell lines established from three different sgRNAs, four in PANC-1 cells and three in MIA PaCa-2 cells, were chosen for use in the study. Please see table at end of methods for primer sequences of guides used.

### Western blotting

For protein extraction from cells, cells were kept on ice and washed three times with PBS, then scraped into PBS and spun down at 200g for 5 minutes. The cell pellet was resuspended in 50-100 μL RIPA buffer [1% NP-40, 0.5% deoxycholate, 0.1% SDS, 150 mM NaCl, 50 mM Tris plus protease inhibitor cocktail (Sigma-Aldrich, P8340) and phosSTOP (Sigma-Aldrich, 04906845001)] and lysis was allowed to continue on ice for 10 minutes. Cells were sonicated with a Fisherbrand Model 120 Sonic Dismembrator (Fisher Scientific, FB120A110) for three pulses of 20 seconds each at 20% amplitude. Cell lysate was spun down at 15,000g for 10 minutes at 4°C and supernatant was transferred to a new tube. For protein extraction from tissue, the sample was resuspended in 500 μL RIPA buffer and homogenized using a TissueLyser (Qiagen, 85210) twice for 30s at 20 Hz. Following incubation on ice for 10 minutes the same procedure was followed as for cells. For both cells and tissue, lysate samples were stored at −80°C until analysis by immunoblot. All blots were developed using a LI-COR Odyssey CLx system. Antibodies used in this study were: O-GlcNAc CTD110.6 (Cell Signaling 9875S), tubulin (Sigma T6199), HSP60 (Cell Signaling 12165S), NAGK (Atlas Antibodies, HPA035207), and PARP (Cell Signaling 9532).

For blots showing the mobility shift for NAGK in low glutamine, samples were prepared in lysis buffer containing 50 mM Tris pH 8.0, 150 mM NaCl, 0.5% IGEPAL CA-630 (Sigma, I3021), 1 mM PMSF, 1.5 μM aprotinin, 84 μM leupeptin, 1 μM pepstatin A, −/+ 10 mM NaF and 20 mM Na_3_VO_4_ as indicated. To visualize the NAGK mobility shift in response to low glutamine, 20 μg total protein per sample was separated across 12.5 cm of 11% SDS-PAGE resolving space under reducing conditions using the large electrophoresis systems available from C.B.S. Scientific until approximately 3 cm of separation was obtained between the 25 and 37 kDa protein standards (Bio-Rad; 1610375). Using electrophoresis, proteins were transferred (30 V, 4°C, overnight) to 0.45 uM pore size nitrocellulose membrane (Amersham, 10600002). The primary antibodies used were NAGK (Proteintech, 15051-1-AP), AKT phosphoS473 (Cell Signaling Technology, 4060), and Vinculin (Sigma, V9264). Membranes were developed using the LI-COR Odyssey CLx system.

### RT-qPCR

For RNA extraction from cells, cells were put on ice, washed with PBS, and scraped into PBS. Samples were then spun down at 200g for 5 minutes and resuspended in 100 μL Trizol (Life Technologies). For RNA extraction from tissue, samples were resuspended in 500 μL Trizol and homogenized using a TissueLyser twice for 30s at 20 Hz. For both cells and tissue, RNA was extracted following the Trizol manufacturer protocol. cDNA was prepared using high-capacity RNA-to-cDNA master mix (Applied Biosystems, 4368814) according to kit instructions. cDNA was diluted 1:20 and amplified with PowerUp SYBR Green Master Mix (Applied Biosystems, A25778) using a ViiA-7 Real-Time PCR system. Fold change in expression was calculated by the ΔΔC_t_ method using HPRT as a control. Please see table at end of chapter for primer sequences.

### Lectin binding assay

Cells were put on ice, washed with PBS and then scraped into PBS. Samples were then spun down at 200g for 5 minutes and resuspended in 3% BSA with fluorophore-conjugated lectin added 1:1000 (Vector Labs FL-1111-2). Samples were covered and incubated on ice for 30 minutes at room temperature, then spun down and resuspended in PBS before analysis with an Attune NxT Flow Cytometer (Thermo Fisher Scientific). Data was further analyzed using FlowJo 8.7.

### Metabolite quantitation and labeling

For all metabolite quantitation experiments, each sample was collected from a 10 cm subconfluent plate of cells. To achieve similar confluency and protein content at the experiment endpoint, cells were initially plated more densely for the nutrient deprived samples than for the nutrient replete samples. For low glutamine experiments, PANC-1 cells were plated 3×10^5^ for 4 mM glutamine samples and 5.5×10^5^ for 0.05 mM samples. MIA PaCa-2 cells were plated 3×10^5^ for 4 mM samples and 1.2×10^6^ for 0.05 mM samples.

Samples were prepared according to Guo et al. (Guo et al., 2016). Briefly, cells were put on ice and washed 3x with PBS. Then, 1 mL of ice cold 80% methanol was added to the plate, and cells were scraped into solvent and transferred to a 1.5 mL tube. For quantitation experiments, internal standard containing a mix of ^13^C labeled metabolites was added at this time. Samples were then sonicated and spun down, and the supernatants were dried down under nitrogen. The dried samples were then resuspended in 100 μL of 5% sulfosalicylic acid and analyzed by liquid chromatography-high resolution mass spectrometry as reported (Guo et al., 2016) with the only modification that the LC was coupled to a Q Exactive-HF with a heated ESI source operating in negative ion mode alternating full scan and MS/MS modes. The [M-H]^-^ ion of each analyte and its internal standard was quantified, with peak confirmation by MS/MS. GlcNAc quantification was done on a triple quadropole instrument exactly as described (Guo et al., 2016). Data analysis was conducted in Thermo XCalibur 3.0 Quan Browser and FluxFix (Trefely et al., 2016). For quantitation experiments, samples were normalized first to peak integrations of ^13^C-labeled internal standard components and then to protein content in the sample, measured by BCA assay. Relative quantification was then calculated by normalizing to the control condition in each experiment.

For glucose labeling experiments, cells were cultured in DMEM without glucose, glutamine, or phenol red supplemented with 10 mM [U-^13^C]-glucose (Cambridge Isotopes, CLM-1396-1), 4 mM glutamine, and 10% dialyzed fetal bovine serum. Cells were incubated for the indicated time, and samples were prepared as above. For GlcNAc labeling experiments, cells were cultured in DMEM without glucose, glutamine, or phenol red supplemented with 10 mM N-[1,2-13C2]acetyl-D-glucosamine (^13^C GlcNAc) (Omicron Biochemicals, GLC-006), 4 mM glutamine, 10 mM glucose, and 10% dialyzed fetal bovine serum. Cells were incubated for the indicated time, and samples were prepared as above.

### Soft agar colony formation assay

Cells were trypsinized and counted using a Bright-Line hemacytometer (Sigma, Z359629). The bottom agar layer was prepared by adding Bacto Agar (BD Bioscience, 214050) to cell culture media for a final concentration of 0.6%. 2 mL bottom agar was added to each well of a 6-well tissue culture plate. Once bottom agar solidified, top layer agar was prepared by combining trypsinized cells with the bottom agar mix for a final concentration of 0.3% Bacto Agar. 1 mL top layer agar was added to each well with a bottom layer of agar. Cells were plated 2.5×10^4^ per well. 0.5 mL DMEM high glucose with 10% calf serum was added to cells every 7 days. Images were taken after 3 weeks. Images were blinded and colonies per image were counted using ImageJ (Schneider et al., 2012).

### 2D Proliferation assay

Cells were plated 3.5×10^4^ per well of a 6-well plate. For each day that counts were recorded, three wells were trypsinized and cells were counted twice using a hemocytometer (Sigma, Z359629). The average of the two counts was recorded for each well, and the average count of the three wells was used to graph the data. For proliferation assays in 0.05 mM glutamine, trypan blue was used during cell counts.

### Bioinformatics data analysis

The PDAC expression profiling dataset (GEO accession GSE16515, Pei et al., 2009) from NCBI GEO Profile database (Edgar et al., 2002) was used to compare the expression level between human normal and PDAC tumor samples. The dataset consists of 52 samples, in which 16 samples are matched tumor and normal tissues, and 20 samples are only tumor tissues. The statistical analysis was conducted by one-way ANOVA, the level of significance was evaluated by p < 0.01 and plotted in box-and-whisker diagram. Comparison of HBP gene expression between tumor (TCGA PAAD dataset) and normal tissue (GTEx) was also conducted using GEPIA2 (Tang et al., 2019).

### Tumor growth in vivo

3×10^6^ PANC-1 NAGK CRISPR cells were injected with 1:1 Matrigel (Corning, CB354248) into the flanks of NCr nude mice and measured with calipers once per week for 22 weeks. At the experiment endpoint (22 weeks or when tumor reached 20 mm in length), mice were euthanized with CO_2_ and cervical dislocation. Tumors were removed, weighed, cut into pieces for analysis, and frozen. All animal experiments were approved by the University of Pennsylvania and the Institutional Animal Care and Use Committee (IACUC).

## Results

### Tetra-antennary N-glycans and O-GlcNAcylation are minimally impacted by nutrient limitation in pancreatic cancer cells

To examine the effects of nutrient deprivation on glycosylation, we cultured cells under glucose or glutamine limitation and examined O-GlcNAc levels and cell surface phytohemagglutinin-L (L-PHA) binding, a readout of N-acetylglucosaminyltransferase 5 (MGAT5)-mediated cell surface N-glycans (Fig S1A, B), which are highly sensitive to UDP-GlcNAc availability (Lau et al., 2007). We focused on glucose and glutamine because of their requirement to initiate the HBP (Fig. 1A). First, as a positive control, we examined HCT-116 and SW480 colon cancer cells, previously documented to have glucose-responsive O-GlcNAcylation (Park et al., 2010; Steenackers et al., 2016), which we also confirmed in HCT-116 cells (Fig. 1B). Indeed, L-PHA binding was suppressed by glucose restriction in SW480 cells and by glutamine restriction in both colon cancer cell lines (Fig. 1C). Next, to test whether glycans were sensitive to nutrient restriction in PDA cells, we examined L-PHA binding and O-GlcNAc levels under nutrient deprivation conditions in a panel of human PDA cell lines, including PANC-1, MIA PaCa-2, AsPC-1, and HPAC. Across these cell lines, no consistent changes in L-PHA binding were observed under glucose or glutamine limitation (Fig. 1D, E; Fig. S1C). We also examined L-PHA binding in PDA cells under oxygen- or serum-deprived conditions and observed minimal changes (Figure S1D, E). O-GlcNAcylation was minimally altered by culture in low glutamine and exhibited variable changes in response to glucose limitation (Fig. 1F), consistent with stress-induced regulation of this modification (Taylor et al., 2008). Since glycosylation may be maintained through either sustained ability to add the modifications or through changes in turnover, we assayed active O-GlcNAcylation by inhibiting O-GlcNAcase with Thiamet G (TMG). TMG treatment resulted in equivalently elevated O-GlcNAcylation levels in high and low glutamine conditions (Fig. 1G; Fig. S1F), indicating that glutamine restriction does not limit the capacity of cells to add the O-GlcNAc modification. Mia-PaCa-2 cells exhibited some cell death in low glutamine, though this was not exacerbated by TMG treatment (Fig. S1F). Thus, under a variety of nutrient stress conditions, neither L-PHA binding nor O-GlcNAcylation were consistently suppressed in pancreatic cancer cell lines. Glutamine restriction in particular had remarkably little impact on O-GlcNAcylation and L-PHA binding, raising the question of how UDP-GlcNAc is generated during nutrient limitation.

**Figure 1:**
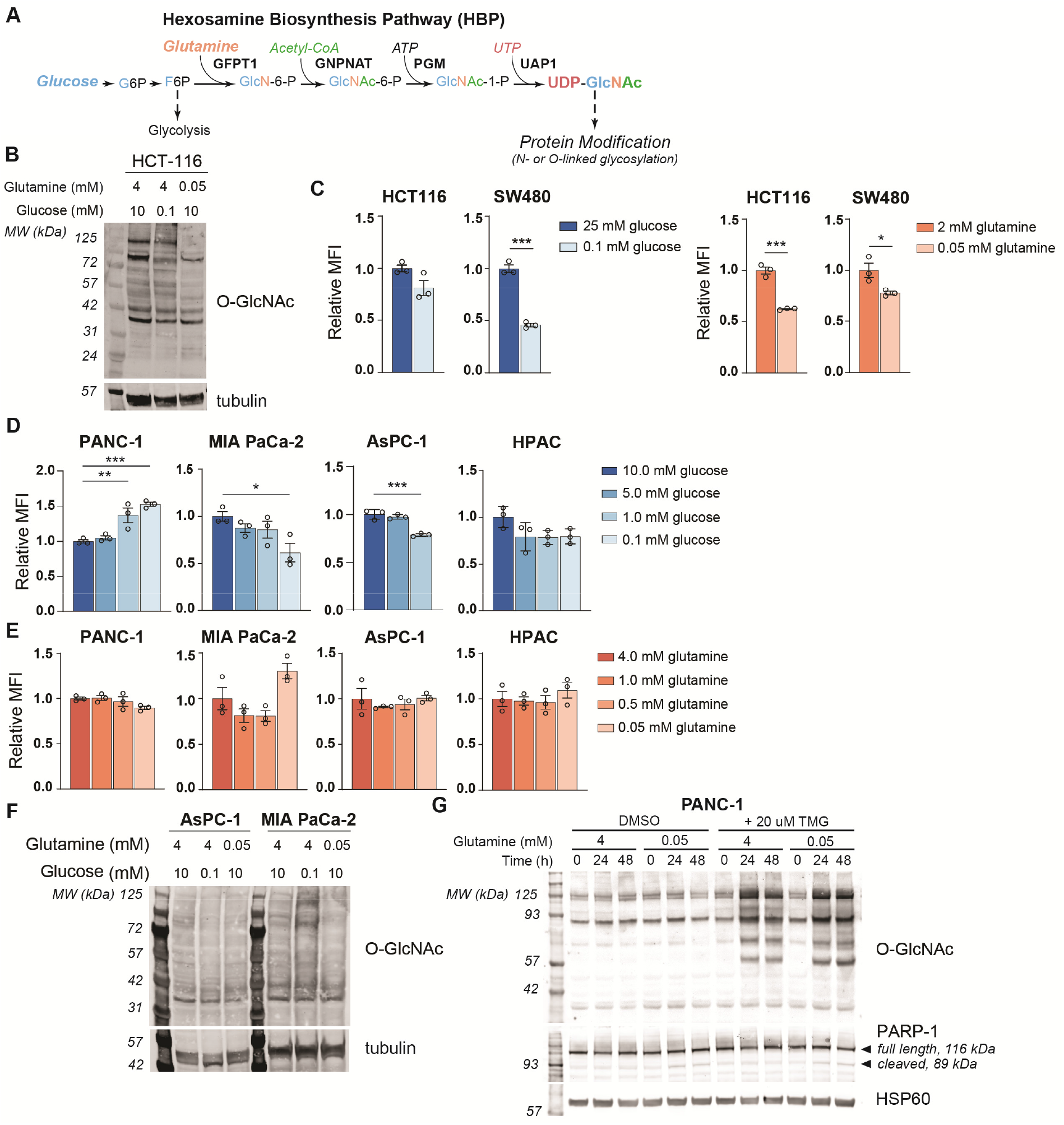
MGAT5-dependent N-glycans are minimally impacted by glucose or glutamine deprivation in PDA cells. **A)** Overview of the hexosamine biosynthesis pathway (HBP). **B)** O-GlcNAc levels in HCT-116 cells in high and low nutrients; cells were incubated in indicated ∞ncentrations of glucose and glutamine for 48 hours. **C)** Phytohemagglutinin-L (LPHA) binding in colon cancer cells. Cells were incubated in the indicated concentrations of glucose (left) or glutamine (right) for 48 hours and then analyzed by flow cytometry. Graph shows mean fluorescence intensity (MFI) relative to control condition. Statistical significance was calculated by unpaired t-test. **D-E)** Phytohemagglutinin-L (LPHA) binding in pancreatic ductal adenocarcinoma (PDA) cells in low nutrients. Cells were incubated in the indicated concentrations of glucose (D) or glutamine (E) for 48 hours and then analyzed by flow cytometry. Statistical significance was calculated by one-way ANOVA. **F)** O-GlcNAc levels in PDA cells in high and low nutrients. Cells were incubated in the indicated concentrations of glu∞se or glutamine for 48 hours. **G)** Western blot for O-GlcNAc and PARP in PANC-1 cells cultured in the indicated concentrations of glutamine with or without Thiamet-G (TMG) treatment for the indicated time. For all bar graphs, mean +/- standard error of the mean (SEM) of three biological triplicates is represented. Panels B) – G) are representative of at least two independent experimental replicates. *, p ≤ 0.05; “, p ≤ 0.01; ***, p ≤ 0.001.

### De novo UDP-GlcNAc synthesis is suppressed upon glutamine limitation

We therefore next asked whether the abundance of HBP metabolites is impacted by nutrient limitation. We measured HBP metabolites after glucose or glutamine restriction using HPLC-MS (Guo et al., 2016). In low glutamine conditions, GlcN-6-P levels were potently decreased relative to glutamine replete conditions in PANC-1 cells, while UDP-GlcNAc abundance was maintained (Fig. 2A). In MIA PaCa-2 cells, UDP-GlcNAc abundance actually increased upon glutamine restriction (Fig S2.1A). These data indicate that UDP-GlcNAc might be generated through mechanisms other than de novo synthesis. Glycolytic intermediates were minimally impacted by low glutamine conditions, and TCA cycle intermediates such as α-KG and malate decreased as expected (Fig. 2A, Fig. S2.1A). In contrast to that in glutamine restriction, UDP-GlcNAc abundance declined in 5 mM or 0.1 mM relative to 10 mM glucose conditions (Fig. S2.1B), suggesting that glutamine limitation specifically may trigger an adaptive response to sustain UDP-GlcNAc pools.

**Figure 2:**
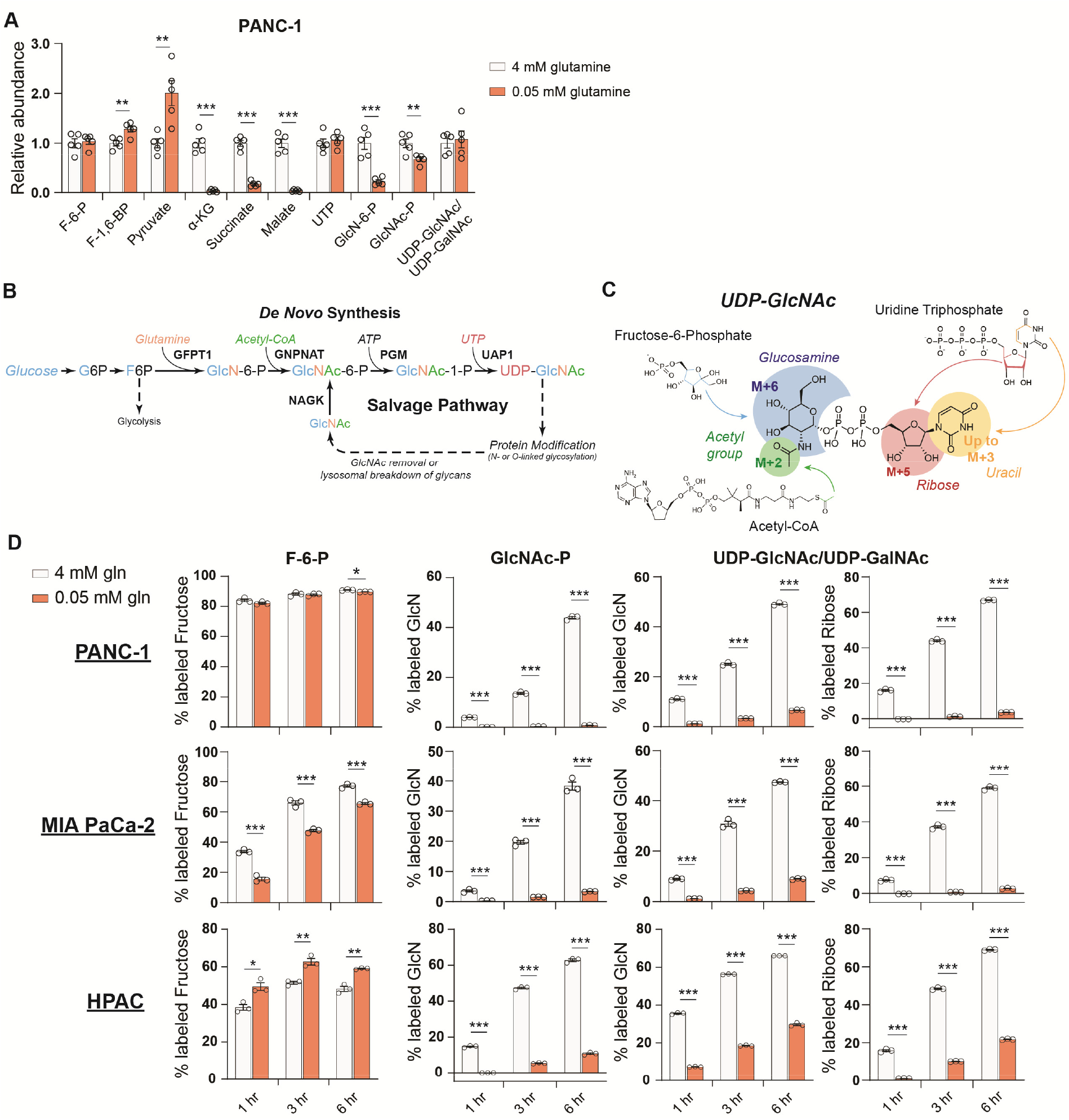
*De novo* UDP-GlcNAc synthesis is supressed upon glutamine limitation. **A)** Metabolite measurements in PANC-1 and MIA PaCa-2 cells after culture for 48 hours in 0.05 mM glutamine. Quantification is normalized to 4 mM glutamine condition. Statistical significance was calculated by unpaired t-test.Mean +/- SEM is represented. **B)** Overview of the GlcNAc salvage pathway feeding into the HBP. GlcNAc scavenged from O-GlcNAc removal or lysosomal breakdown of glycans can be phosphorylated by NAGK and used to regenerate UDP-GlcNAc. **C)** Overview of the in∞rporation of ^13^C glu∞se into UDP-GlcNAc. Different parts of the molecule can be labeled from glucose-derived subunits, thus isotopologues up to M+16 can be derived from glucose. **D)** ^13^C glucose tracing into F-6-P, GlcNAc-P, and UDP-GlcNAc in indicated glutamine concentrations. % labeled GlcN indicates sum of M+6 and M+8 isotopologues for GlcNAc-P and sum of M+6, M+8, M+11, and M+13 for UDP-GlcNAc, % labeled Ribose indicates sum of M+5, M+7, M+11, and M+13 for UDP-GlcNAc. All isotopologues are graphed in Figure S2. Statistical significance was calculated by unpaired t-test. Mean +/- SEM is represented. *, p ≤ 0.05; **, p ≤ 0.01; ***, p ≤ 0.001.

We sought to understand how UDP-GlcNAc pools are sustained during glutamine restriction. In addition to de novo synthesis of UDP-GlcNAc via the HBP, free GlcNAc in the cell can also be phosphorylated via N-acetylglucosamine kinase (NAGK) to produce GlcNAc-P and then regenerate UDP-GlcNAc (Fig. 2B). However, NAGK’s roles in physiology and cancer biology have been minimally studied. To investigate the possibility that UDP-GlcNAc is generated through mechanisms other than its synthesis from glucose, we first designed a stable isotope labeling strategy to quantify the fraction of the glucosamine ring that is synthesized *de novo* in glutamine-replete versus -restricted conditions. Since multiple components of UDP-GlcNAc [glucosamine ring, acetyl group, uridine (both the uracil nucleobase and the ribose ring)] can be synthesized from glucose, UDP-GlcNAc isotopologues up to M+16 can be generated from glucose (Moseley et al., 2011) (Fig. 2C). In order to measure the glucose carbon incorporated into GlcNAc-P and UDP-GlcNAc via the HBP, all isotopologues containing a fully labeled glucosamine ring are added together (% labeled GlcN indicates sum of M+6, M+8, M+11, and M+13 for UDP-GlcNAc and sum of M+6 and M+8 for GlcNAc-P) (Fig. 2C).

After 48 hours of glutamine restriction, cells were incubated with fresh low glutamine medium containing [U-^13^C]-glucose to track the incorporation of glucose carbons into hexosamine intermediates. Across multiple PDA cell lines, the fractional labeling of the glucosamine ring in both GlcNAc-P and UDP-GlcNAc pools was markedly suppressed by glutamine restriction, indicating decreased de novo synthesis in low glutamine conditions (Fig. 2D, Fig. S2.1C-E). Notably, labeling into the ribose component of UDP-GlcNAc was also suppressed [% labeled ribose indicates sum of isotopologues containing M+5 (i.e., M+5, M+7, M+11, and M+13); Fig. 2D, Fig. S2.1C]. Consistently, incorporation of ^13^C glucose into uridine triphosphate (UTP) was suppressed upon glutamine restriction (Fig. S2.2A), even though UTP levels were maintained or increased (Fig. 2A, Fig. S2.1A), suggesting a role for nucleoside salvage in maintaining nucleotide pool in these conditions. This is consistent with previous reports demonstrating that autophagy/ ribophagy is a source of nucleosides in amino acid deprived conditions (Guo et al., 2011; Wyant et al., 2018). Indeed, silencing of either of the uridine salvage enzymes uridine kinase 1 or 2 (UCK1/2) resulted in decreased UDP, UTP, and UDP-GlcNAc levels (Fig. S2.2B-C), indicating that nucleoside salvage contributes to maintaining uridine phosphate and UDP-GlcNAc pools.Thus, glutamine restriction suppresses the *de novo* synthesis of both GlcNAc-P and UTP, both of which are required to produce UDP-GlcNAc.

We noted that GlcNAc-P and UDP-GlcNAc pools labeled from glucose with similar but not identical kinetics. While this is potentially due to limitations in detection since GlcNAc-P is much less abundant than UDP-GlcNAc, we considered whether GlcNAc-P-independent pathways may also have minor contributions to glucose-dependent UDP-GlcNAc labeling. Although pathways through which glucose can feed into UDP-GlcNAc’s glucosamine ring independent of the HBP have not been described in mammalian cells to our knowledge, we nevertheless tested the two major metabolic branch points diverging from UDP-GlcNAc, which mediate UDP-GalNAc and sialic acid synthesis. UDP-galactose-4-epimerase (GALE) interconverts UDP-GlcNAc and UDP-GalNAc, and UDP-GlcNAc-2-epimerase/ManAc kinase (GNE) initiates sialic acid biosynthesis. Silencing of neither GALE nor GNE reduced UDP-GlcNAc labeling from glucose, however, indicating that these enzymes are unlikely to facilitate a bypass pathway (Fig. S2.2D-E). Although a minor contribution from another unknown pathway cannot be ruled out, the slight apparent differences in timing of GlcNAc-P and UDP-GlcNAc labeling most likely reflect technical limitations. Regardless, the data clearly indicate that UDP-GlcNAc abundance is maintained despite reduced *de novo* hexosamine synthesis from glucose (Fig. 2D).

### GlcNAc salvage feeds UDP-GlcNAc pools in pancreatic cancer cells

As mentioned, UDP-GlcNAc can be generated via phosphorylation of free GlcNAc by NAGK generating GlcNAc-6-P (Fig. 2B). When supplemented, GlcNAc is salvaged into the UDP-GlcNAc pool (Ryczko et al., 2016; Wellen et al., 2010). Endogenous sources of GlcNAc may include removal of O-GlcNAc protein modifications or breakdown of glycoconjugates and extracellular matrix components. Notably, intracellular levels of GlcNAc increase upon glutamine restriction (Fig. 3A). Yet, the significance of GlcNAc salvage to maintenance of UDP-GlcNAc pools has been little studied, and the proportion of UDP-GlcNAc generated via the NAGK-dependent salvage pathway is unknown.

**Figure 3:**
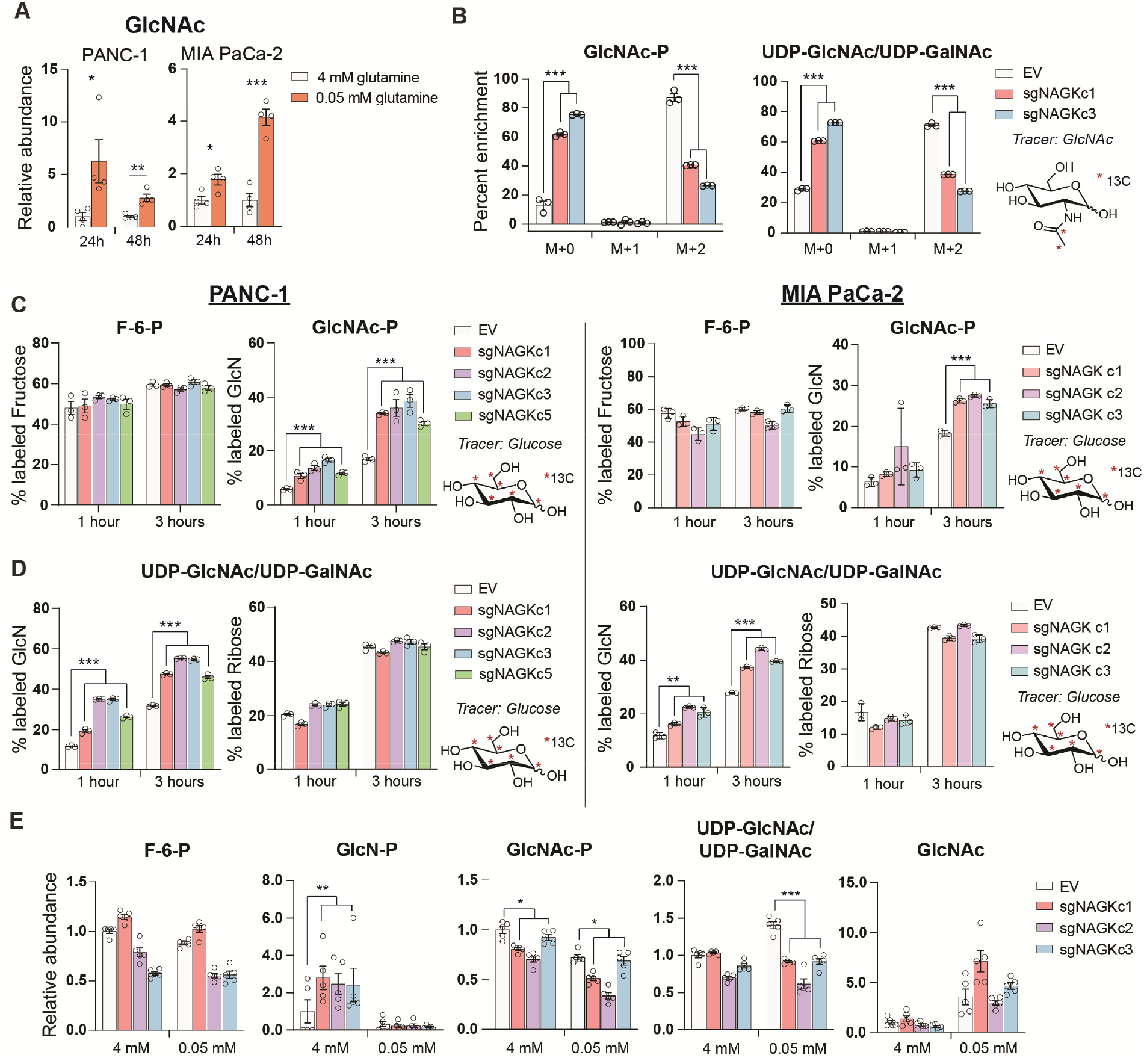
GlcNAc salvage feeds UDP-GlcNAc pools in pancreatic cancer cells. **A)** Measurement of GlcNAc in PANC-1 and MIA PaCa-2 cells after incubation in the indicated concentrations of glutamine for 24 and 48 hours. Mean +/- SEM of four biological replicates is represented. Statistical significance was calculated by unpaired t-test. **B)** Measurement of 13C GlcNAc labeled on the acetyl group into GlcNAc-P and UDP-GlcNAc in NAGK knockout cells. Cells were incubated with 10 mM 13C GlcNAc for 6 hours. Mean +/- SEM of three biological replicates is represented. Statistical significance was calculated by unpaired t-test comparing the mean incorporation of the two CRISPR clones and the empty vector (EV) control. **C)** Labeling of F-6-P and GlcNAc-P from 13C glucose in PANC-1 (left) and MIA PaCa-2 (right) NAGK knockout cells. Statistical significance was calculated by unpaired t-test comparing the mean incorporation of the four CRISPR clones and the EV control. Mean +/- SEM of three biological replicates is represented. **D)** Percent of combined UDP-GlcNAc isotopologues containing a labeled glucosamine ring or a labeled ribose from UTP, calculated from S3.1 (E). Statistical significance was calculated by unpaired t-test comparing the mean incorporation of the four CRISPR clones and the EV control. Mean +/- SEM of three biological replicates is represented. **E)** Measurement of HBP metabolites in PANC-1 NAGK knockout cells cultured in the indicated concentrations of glutamine. Statistical significance was calculated by unpaired t-test comparing the mean incorporation of the four CRISPR clones and the EV control. Mean +/- SEM of four biological replicates is represented. *, p ≤ 0.05; **, p ≤ 0.01; ***, p ≤ 0.01.

*NAGK* mRNA expression increased in PDA cell lines in low glutamine conditions, and in some cell lines also in low glucose (Fig. S2.3A, B). *GFPT1* expression was also induced in both low glucose conditions, consistent with a prior report (Moloughney et al., 2016), and in low glutamine conditions (Fig. S2.3A), even though *de novo* synthesis is suppressed when glutamine is limited. Protein levels of NAGK did not increase in concordance with mRNA at these time points, however, although a mobility shift potentially indicative of post-translational modification was apparent when protein lysates were run on a gel using a large electrophoresis system (see methods; Fig S2.3C-D). Removal of the phosphatase inhibitor Na_3_VO_4_ from the sample buffer prevented the mobility shift, suggesting that NAGK may be phosphorylated on one or more residues in low glutamine conditions (Fig. S2.3D). Taken together, these data indicate that under low glutamine conditions, GlcNAc availability for salvage increases and the salvage enzyme NAGK is subject to regulation.

These findings prompted us to investigate the role of NAGK in UDP-GlcNAc synthesis in PDA cells. We functionally examined the role of NAGK in PDA cell lines by using CRISPR-Cas9 gene editing to generate NAGK knockout (KO) PANC-1 and MiaPaCa-2 clonal cell lines (Fig. S3.1A-B). N-[1,2-^13^C_2_]acetyl-D-glucosamine (^13^C GlcNAc) was efficiently salvaged in control cells, and this was suppressed by NAGK deletion, as evidenced by reduced fractional labeling of GlcNAc-P and UDP-GlcNAc (Fig. 3B). Since we did not observe any residual protein expression, we hypothesized that the N-acetylgalactosamine (GalNAc) salvage enzyme GalNAc kinase (GALK2) might be responsible for the remaining GlcNAc salvage in the absence of NAGK. Indeed, silencing of GALK2 further suppressed incorporation of ^13^C GlcNAc into GlcNAc-P and UDP-GlcNAc in the NAGK KO cells (Fig. S3.1C).

We hypothesized that knockout cells would conversely conduct increased de novo UDP-GlcNAc synthesis. To test this, we incubated cells with [U-^13^C]-glucose and examined incorporation into GlcNAc-P and UDP-GlcNAc. Indeed, in the absence of NAGK, we observed increased glucose-dependent fractional labeling of the glucosamine ring of UDP-GlcNAc and GlcNAc-P, but not the ribose component of UDP-GlcNAc (Fig. 3C-D; Fig. S3.1D-E). This effect was also observed with knockdown of NAGK by shRNA, though to a lesser extent (Fig. S3.2A-B). Incorporation of glucose into F-6-P did not change (Fig. 3C) and the proportion of UDP-GlcNAc containing an M+5 ribose ring was also unchanged in knockout cells (Fig. 3D), as expected. Thus, when NAGK is deleted and GlcNAc salvage is suppressed, de novo hexosamine synthesis increases.

We next assessed changes in the levels of hexosamine intermediates in control and NAGK KO cell lines. In PANC-1 KO cells in 4 mM glutamine, GlcN-P increased significantly, consistent with increased de novo synthesis in the absence of NAGK (Fig. 3E). GlcNAc-P was modestly reduced in KO cells, though UDP-GlcNAc levels were maintained (Fig. 3E). In MIA PaCa-2 cells, GlcNAc-P was markedly suppressed in the absence of NAGK, though UDP-GlcNAc was not (Fig. S3.2C). We also measured HBP metabolites under glutamine restriction, where we expected NAGK would play a more significant role in UDP-GlcNAc generation. We were only able to measure metabolites accurately in PANC-1 KO cells because MIA PaCa-2 NAGK KO cells began to die quickly in low glutamine, which will be discussed further in the next section. GlcN-P decreased in control and KO cells, consistent with reduced de novo hexosamine synthesis (Fig. 3E). GlcNAc-P levels decreased in low glutamine in control cells, and decreased further in cells lacking NAGK, consistent with contributions from both de novo synthesis and salvage (Fig. 3E). Reciprocally, GlcNAc abundance was elevated upon glutamine limitation in both control and NAGK KO cells (Fig. 3E). UDP-GlcNAc abundance was modestly reduced in NAGK KO cells relative to controls under glutamine restriction, though levels were still comparable to that in high glutamine (Fig. 3E), possibly reflecting changes in utilization. GALK2 silencing did not further suppress UDP-GlcNAc in NAGK KO cells (Fig. S3.2D), suggesting that GALK2 may not have a major role in physiological GlcNAc salvage. Cumulatively, the data demonstrate that GlcNAc is salvaged into UDP-GlcNAc pools in PDA cells in a manner dependent at least in part on NAGK.

### NAGK knockout limits tumor growth *in vivo*

To test the role of NAGK in cell proliferation, we first monitored growth of NAGK KO cells compared to controls in 2D and 3D culture in 4 mM glutamine, finding minimal differences (Fig. 4A, Fig. S4A). We hypothesized that NAGK KO cell proliferation would be impaired in 0.05 mM glutamine, where *de novo* UDP-GlcNAc synthesis is suppressed. Indeed, MIA PaCa-2 KO cells died more quickly in 0.05 mM glutamine than did control cells (Fig. 4A). PANC-1 KO cells did not show this effect (Fig. 4A), but we hypothesized that NAGK loss might have a stronger effect *in vivo* where tumor growth can be constrained by nutrient availability.

**Figure 4:**
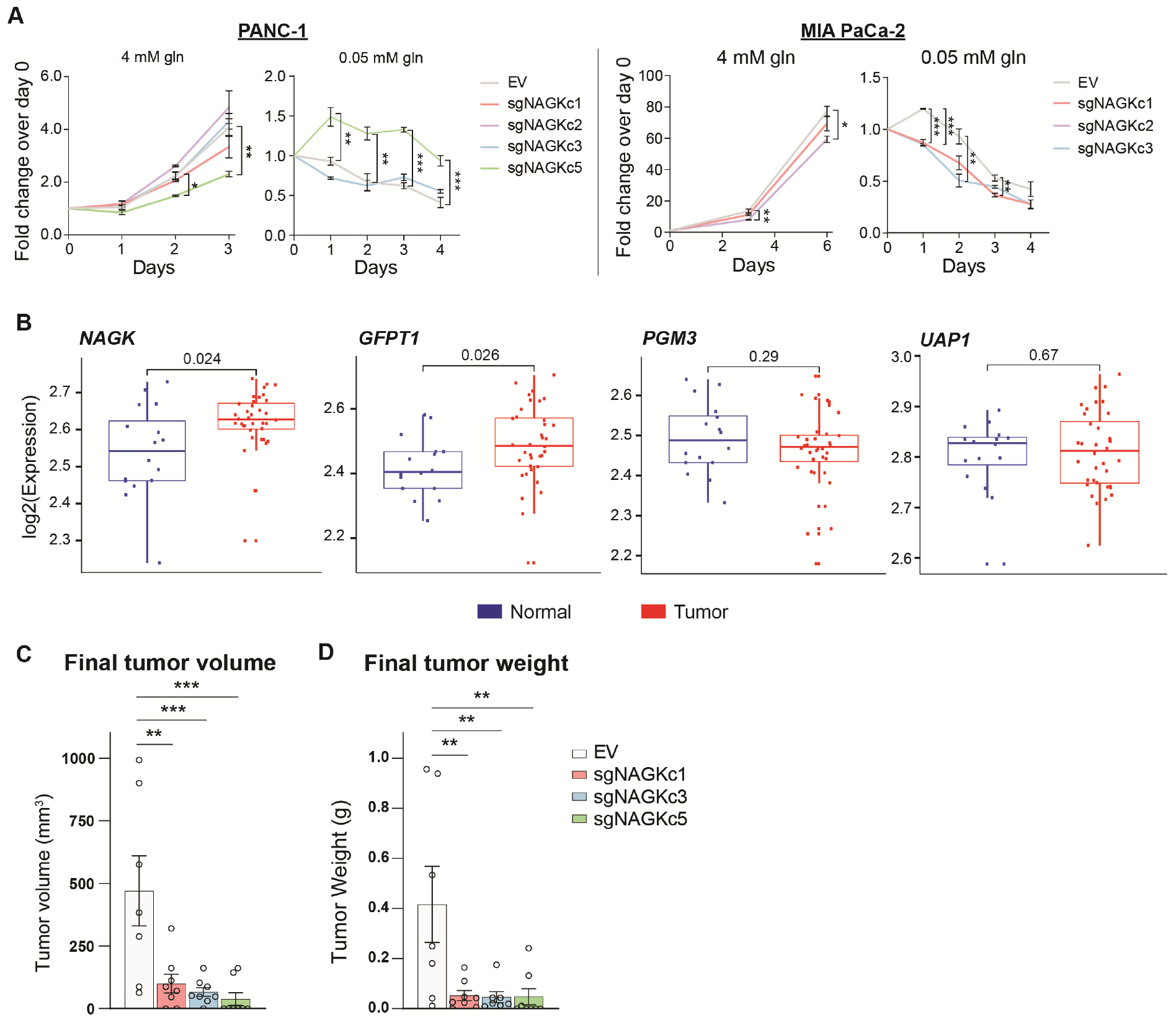
NAGK expression is increased in human PDA tumors and NAGK knockout reduces tumor growth in vivo. **A)** 2D proliferation assay by cell count in PANC-1 and MIA PaCa-2 NAGK knockout cells. Mean +/- SEM of three technical replicates is represented. Statistical significance was calculated using one-way AN OVA at each time point. **B)** Gene expression data for NAGK, GFPT1, PGM3, and UAP1 in human PDA tumors compared with matched normal tissue. Statistical analysis was conducted by one-way ANOVA, and level of significance was defined as p ≤ 0.01. **C)** Final tumor volume and **D)** final tumor weight of subcutaneous tumors generated from PANC-1 NAGK knockout cells in vivo. Cells were injected into the right flank of NCr nude mice and tumor volume was calculated from caliper measurements. Statistical significance was calculated by one-way ANOVAcomparing each mean to the EV control mean. Mean +/- SEM of biological replicates is represented (n = 8 each group). *, p ≤ 0.05; **, p ≤ 0.01; ***, p ≤ 0.001.

To gain initial insight into whether NAGK is likely to play a functional role in PDA progression *in vivo*, we queried publicly available datasets. From analysis of publicly available microarray data (Pei et al., 2009) and gene expression data from the Cancer Genome Atlas (TCGA), we indeed found *NAGK* expression to be increased in tumor tissue relative to adjacent normal regions of the pancreas or to pancreas GTEx data (Fig. 4B, Fig. S4B). *GFPT1* expression was also increased in tumor tissue (Fig. 4B, Fig. S4B), consistent with its regulation by mutant KRAS (Ying et al., 2012). Two other HBP genes, *PGM3* and *UAP1,* did not show significantly increased expression in PDA tumors in these datasets (Fig. 4B, Fig. S4B). We then studied the role of NAGK in tumor growth *in vivo* by injecting NAGK CRISPR KO cells into the flank of NCr nude mice. Final tumor volume and weight were markedly reduced in the absence of NAGK (Fig. 4C-D). Of note, initial tumor growth was comparable between control and KO cells, but the NAGK knockout tumors either stopped growing or shrank while control tumors continued to grow larger (Fig. S4C). Interestingly, KO tumor samples showed increased L-PHA signal (Fig. S4D), indicating that NAGK deficiency results in altered glycosylation within tumors. This could possibly reflect either elevated de novo synthesis in the small tumors that form or differences in cellular composition. For example, activated fibroblast marker α-smooth muscle actin (α-SMA) was more abundant in the NAGK KO tumors (Fig S4D). Residual NAGK signal in whole tumors also presumably reflects expression in other cell types (Fig. S4D), since NAGK was undetectable in the clonal cell lines used for injections (Fig. S3.1B). Taken together, these data are consistent with the notion that NAGK is dispensable when nutrients are abundant but becomes more important as the tumors outgrow their original nutrient supply and become more dependent on scavenging and recycling and indicate that NAGK-mediated hexosamine salvage supports tumor growth *in vivo.*

## DISCUSSION

In this study, we identify a key role for NAGK in salvaging GlcNAc for UDP-GlcNAc synthesis in PDA cells. We show that glutamine deprivation suppresses *de novo* hexosamine biosynthesis, which is reciprocally increased upon NAGK deletion. Glutamine deprivation also results in increased availability of GlcNAc for salvage. *NAGK* expression is elevated in human PDA tumors, and NAGK deficiency suppresses GlcNAc salvage in cells and tumor growth in mice.

This work raises several key questions for future investigation. First, the sources of GlcNAc salvaged by NAGK remain to be fully elucidated. GlcNAc may be derived from recycling of GlcNAc following O-GlcNAc removal or breakdown of glycoconjugates. Additionally, GlcNAc may be recovered from the environment. Nutrient scavenging via macropinocytosis is a key feature of PDA (Commisso et al., 2013; Kamphorst et al., 2015). Macropinocytosis has mostly been associated with scavenging of protein to recover amino acids, but lysosomal break down of glycoproteins may also release sugars including GlcNAc. Additionally, ECM components, including hyaluronic acid (HA), which is a polymer of GlcNAc and glucuronic acid disaccharide units, may be additional sources of GlcNAc for salvage in the tumor microenvironment. Indeed, in a manuscript co-submitted with this one, Kim, Halbrook, and colleagues identify HA as a major source of scavenged GlcNAc (Kim et al., 2020). Our manuscript and the Kim, Halbrook et al manuscript together indicate that NAGK may take on a heightened importance in the context of high GlcNAc availability and nutrient deprivation, a situation that is likely to occur within the tumor microenvironment.

Further, the key fates of UDP-GlcNAc that support tumor growth remain to be elucidated. Sufficient UDP-GlcNAc is required for protein glycosylation to maintain homeostasis and prevent ER stress, particularly in a rapidly dividing cell. Additionally, a wide range of cancers exhibit elevated O-GlcNAc, which could contribute to driving pro-tumorigenic transcriptional and signaling programs. In PDA specifically, the glycan CA19-9 is currently used as a biomarker for disease progression and recent studies point to a functional role for CA19-9 in tumorigenesis (Engle et al., 2019). UDP-GlcNAc is also required for HA synthesis, which is present in low amounts in normal pancreas but increases in PanIN lesions and PDA (Provenzano et al., 2012). PDA cells are capable of producing HA in vitro (Mahlbacher et al., 1992). Depletion of fibroblasts in an autochthonous PDA mouse model results in a decrease in collagen I but not HA in the tumor microenvironment, indicating that HA must be generated by another cell type, possibly the tumor cells themselves (Özdemir et al., 2014). Previous studies demonstrated that treatment of PDA with exogenous hyaluronidase can increase vascularization and improve drug delivery to the tumor (Jacobetz et al., 2013; Provenzano et al., 2012), although a phase III clinical trial reported no improvement in overall patient survival when combining pegylated hyaluronidase with nab-paclitaxel plus gemcitabine (Cutsem et al., 2020). Recently, it was shown that inhibiting the HBP by treatment with 6-diazo-5-oxo-l-norleucine (DON) depletes HA and collagen in an orthotopic mouse model. DON treatment also increased CD8 T-cell infiltration into the tumor, sensitizing the tumor to anti-PD1 therapy (Sharma et al., 2020). Thus, targeting the HBP holds promise for improving the efficacy of other therapeutics. The findings of the current study suggest that in addition to de novo hexosamine synthesis, targeting of hexosamine salvage warrants further investigation in terms of potential for therapeutic intervention. Of note, an inhibitor targeting PGM3, which converts GlcNAc-6-P to GlcNAc-1-P and is thus required for both de novo UDP-GlcNAc synthesis and GlcNAc recycling, showed efficacy in treating gemcitabine-resistant patient-derived xenograft PDA models (Ricciardiello et al., 2020), as well as in breast cancer xenografts (Ricciardiello et al., 2018).

Finally, almost nothing is currently known about the role of NAGK and GlcNAc salvage in normal physiology. Even in non-cancerous IL-3-dependent hematopoietic cells, a substantial proportion of the UDP-GlcNAc pool remains unlabeled from ^13^C-glucose (Wellen et al., 2010), suggesting that salvage may contribute to UDP-GlcNAc pools in a variety of cell types. However, while GFPT1 is required for embryonic development in mice, NAGK knockout mouse embryos are viable (Dickinson et al., 2016). NAGK deficiency has not yet been characterized in postnatal or adult mice. Perhaps GlcNAc salvage is dispensable when nutrients are available and cells are not dividing, as in most healthy tissues. However, in a tumor, in which cells are proliferating and nutrients are spread thin, NAGK and GlcNAc salvage may become more important in feeding UDP-GlcNAc pools. Related questions include elucidating the mechanisms regulating NAGK gene expression and putative post-translational modification, as well as understanding the role of GALK2 in hexosamine salvage.

In sum, we report a key role for NAGK in feeding UDP-GlcNAc pools in PDA cells and in supporting xenograft tumor growth. Further investigation will be needed to elucidate the physiological functions of NAGK, as well as the mechanisms through which it supports tumor growth and its potential role in modulating therapeutic responses.

---

**Supplemental Figure 1:**
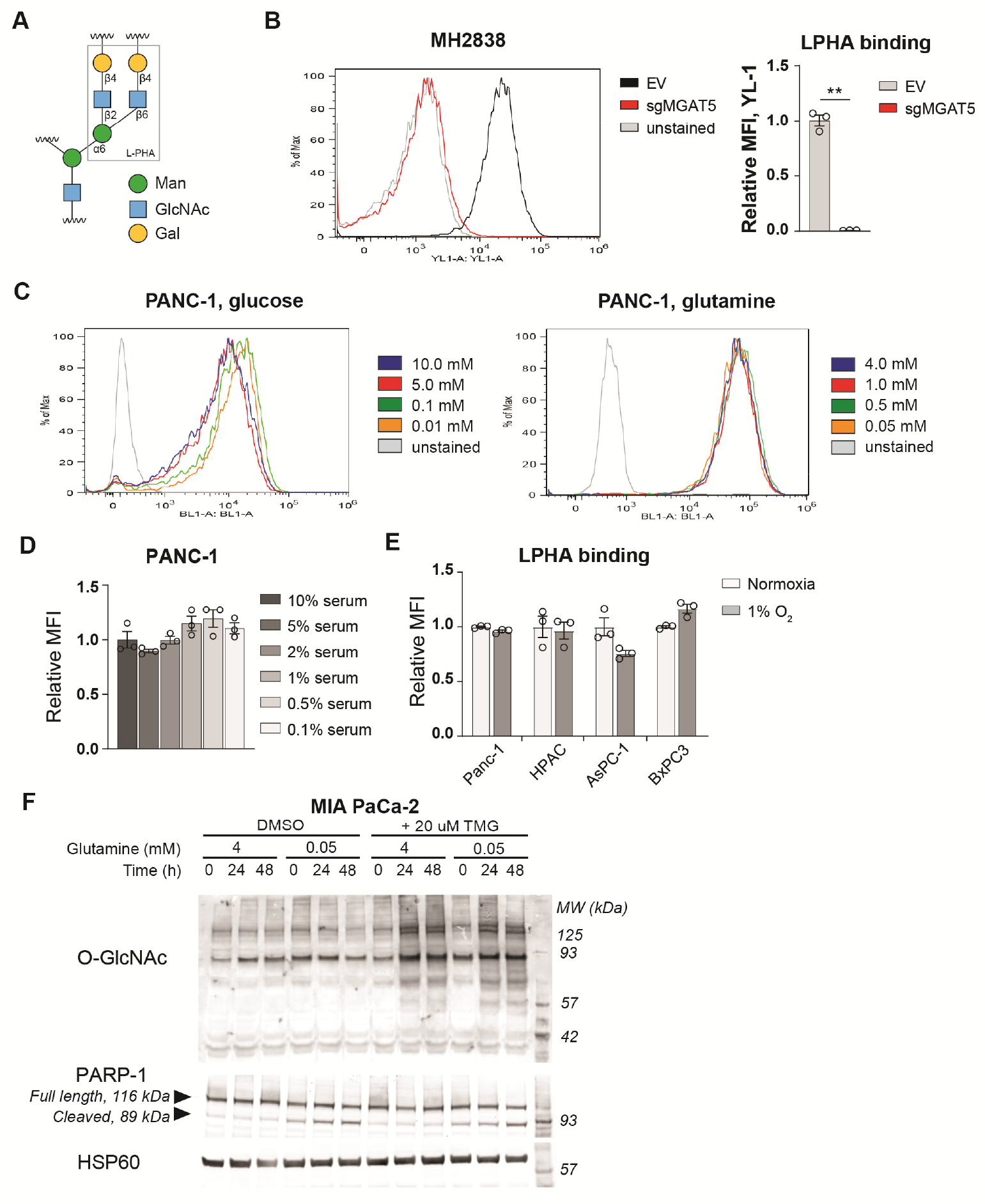
LPHA binding detects MGAT5-dependent glycans. **A)** Diagram of LPHA binding. LPHA recognizes specifically the ß1-6 linkage established by MGAT5. **B)** LPHA binding on MGAT5 knockout cells isolated from a KPCY tumor (Li, Byrne et al., 2018), representative flow plot and quantification. Graph shows MFI relative to control cells. Statistical significance was calculated by unpaired t-test. **C)** Representative flow plots for LPHA binding graphed in Fig. 1D, E on PANC-1 cells in low glucose and low glutamine. **D-E)** LPHA binding in PDA cells in low nutrients. Cells were incubated in the indicated concentrations of serum (D) or oxygen (E) for 48 hours and then analyzed by flow cytometry. Graph shows MFI relative to control condition. **F)** Western blot for O-GlcNAc and PARP in MIA PaCa-2 cells cultured in the indicated concentrations of glutamine with or without TMG for the indicated time. Statistical significance was calculated by one-way ANOVA (D) and unpaired t-test (E). Panels B) – E) are representative of at least two independent experimental replicates. *, p ≤ 0.05; **, p ≤ 0.01; **, p ≤ 0.001.

**Supplemental Figure S2.1:**
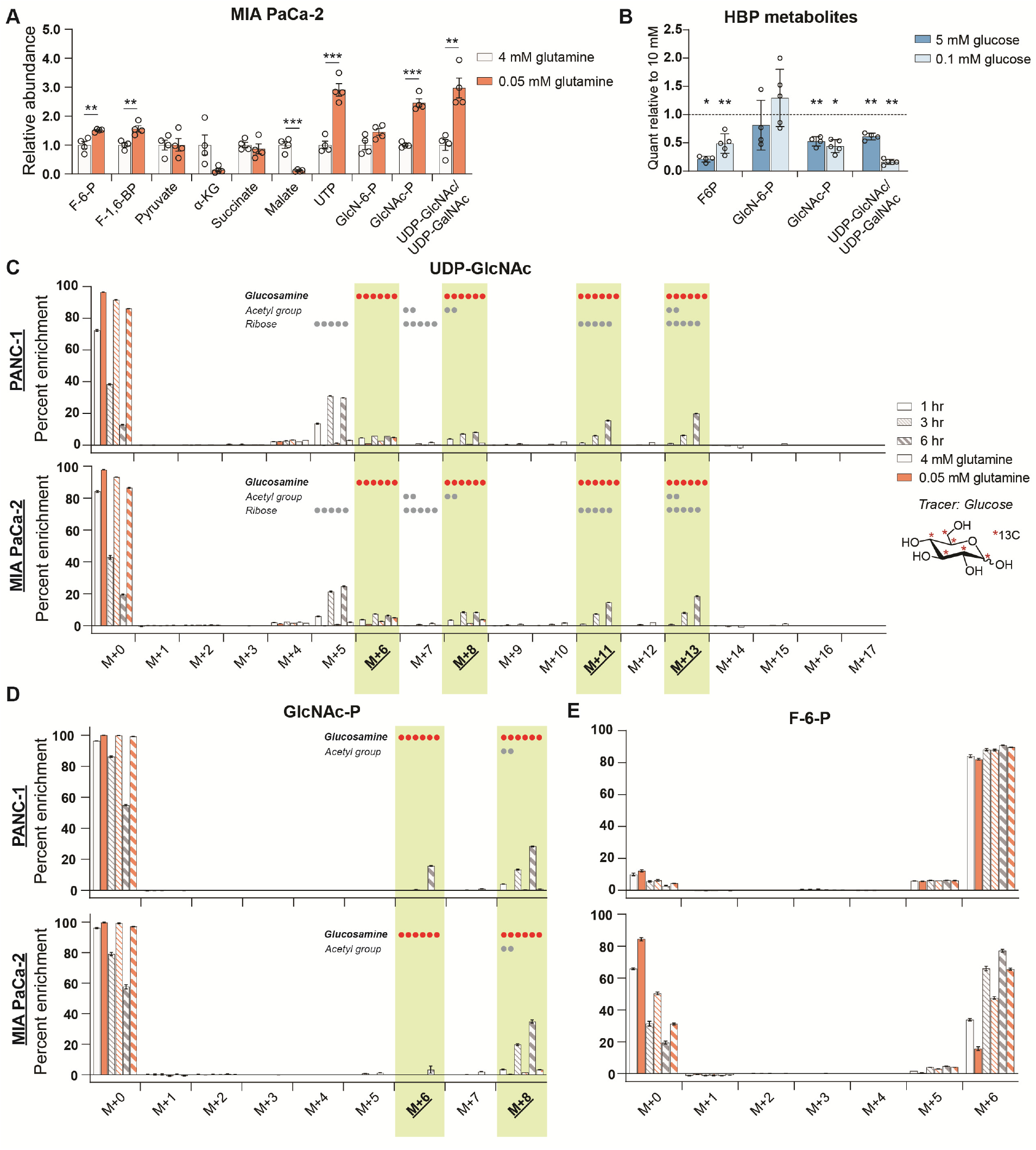
De novo hexosamine synthesis is suppressed in low glutamine conditions. **A)** Metabolite measurements in MIA PaCa-2 cells after culture for 48 hours in 0.05 mM glutamine. Quantification is normalized to 4 mM glutamine condition. Statistical significance was calculated by unpaired t-test. Mean +/- SEM of four biological replicates is represented. **B)** Measurement of HBP metabolites in PANC-1 cells after culture for 48 hours in 5 mM or 0.1 mM glucose. Mean +/- SEM of four (5 mM) or five (0.1 mM) biological replicates is represented. Statistical significance was calculated by unpaired t-test comparing each low glucose condition to the 10 mM glucose control in each experiment. **C)** Measurement of 13C glucose incorporation into UDP-GlcNAc in high and low glutamine. Mean +/- SEM of three biological replicates is represented. **D)** Measurement of 13C glucose incorporation into GlcNAc-P in high and low glutamine. Mean +/- SEM of three biological replicates is represented. For (C-D), components of molecule labeled in each isotopologue are indicated, with dots representing the number of carbons; highlighted panels identify glucosamine ring labeling. **E)** Measurement of 13C glucose into fructose-6-phosphate in high and low glutamine. Mean +/- SEM of three biological replicates is represented. *, p ≤ 0.05; **, p ≤ 0.01; ***, p ≤ 0.001.

**Supplemental Figure S2.2:**
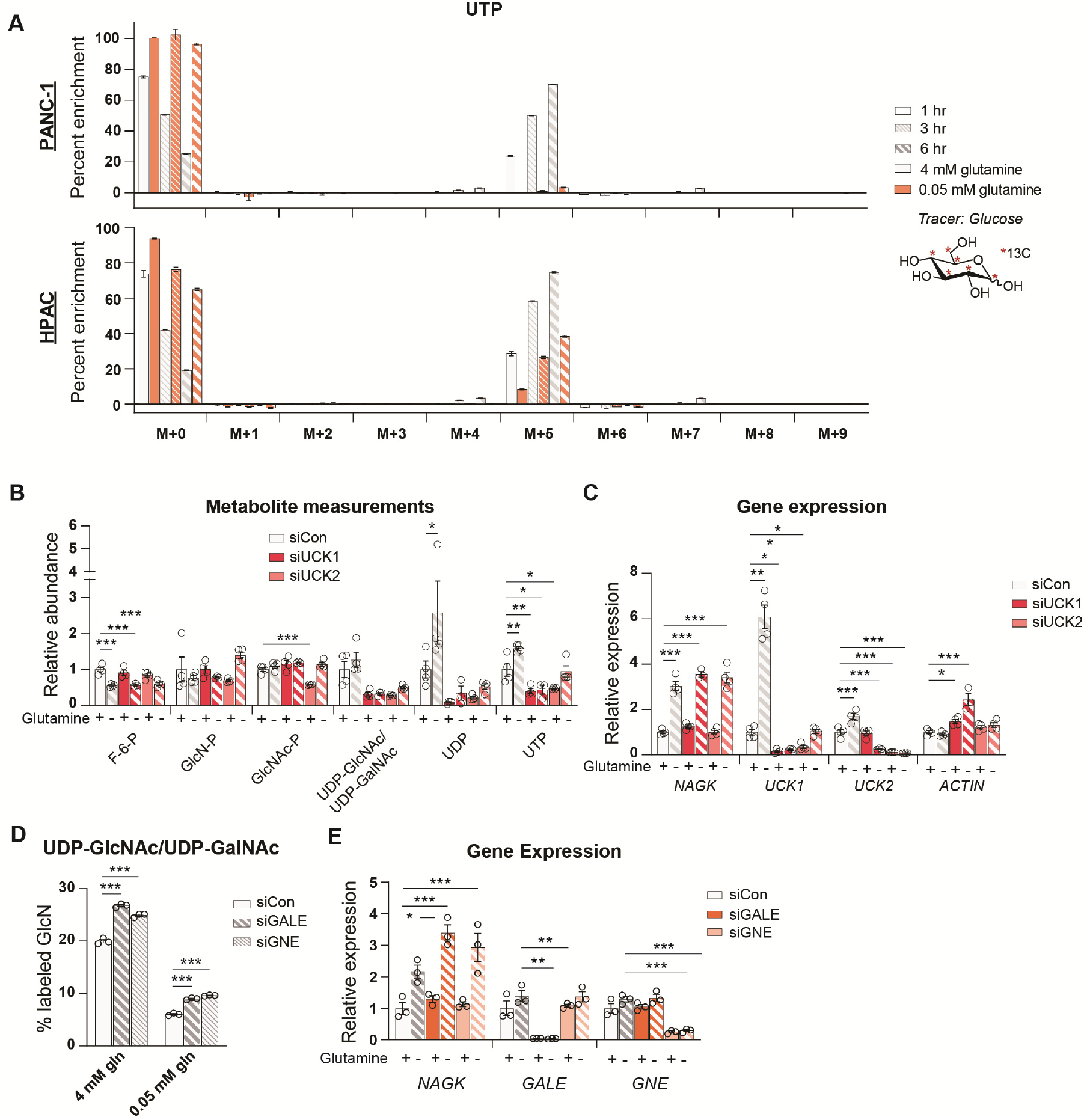
Uridine is salvaged in glutamine restricted conditions. **A)** Measurement of incorporation of 13C glucose into uridine triphosphate (UTP) in high and low glutamine. Mean +/- SEM of three biological replicates is represented. **B)** Measurement of metabolites in PANC-1 cells after transfection with the indicated siRNAs and incubation in 4 mM (+) or 0.05 mM (-) glutamine for 48 hours. Mean +/- SEM of four biological replicates is represented. **C)** Gene expression of the indicated genes in PANC-1 cells after transfection with the indicated siRNAs and incubation in 4 mM (+) or 0.05 mM (-) glutamine for 48 hours. Mean +/- SEM of four biological replicates is represented. **D)** Measurement of incorporation of 13C glucose into UDP-GlcNAc in PANC-1 cells transfected with the indicated siRNAs and incubated in 4 mM or 0.05 mM glutamine for 48 hours. Mean +/- SEM of three biological replicates is represented. **E)** Gene expression of the indicated genes in PANC-1 cells after transfection with the indicated siRNAs and incubation in 4 mM (+) or 0.05 (-)rnM glutamine for 48 hours. Mean +/- SEM of three biological replicates is represented. *, p ≤ 0.05; **, p ≤ 0.01; ***, p ≤ 0.001.

**Supplemental Figure S2.3:**
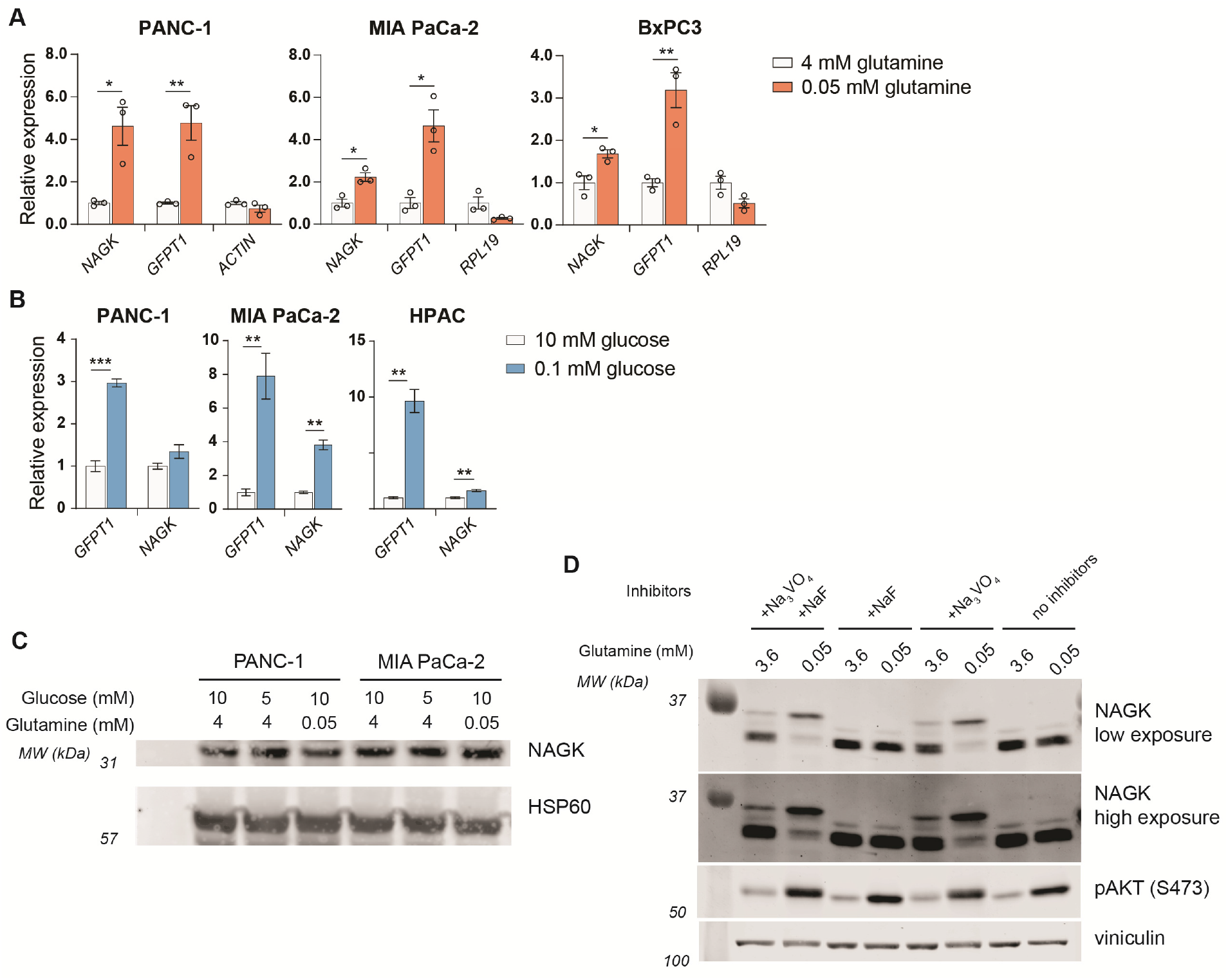
NAGK protein expression does not increase in low glutamine despite increase in NAGK gene expression. **A)** Gene expression of the indicated genes in PDA cell lines after incubation in 4 mM or 0.05 mM glutamine for 48 hours. Mean +/- SEM of three biological replicates is represented. Statistical significance was determined by unpaired t-test. **B)** Gene expression of the indicated genes in PDA cell lines after incubation in 10 mM or 0.1 mM glucose for 48 hours. Mean +/- SEM of three biological replicates is represented. Statistical significance was determined by unpaired t-test. **C)** Western blot for NAGK in low nutrient conditions. PANC-1 and MIA PaCa-2 cells were cultured in the indicated concentrations of glucose and glutamine for 48 hours. **D)** Western blot for NAGK in low nutrient conditions. PANC-1 cells were cultured in the indicated glutamine concentrations and lysates were prepared with or without phosphatase inhibitors. A band shift is observed for NAGK in lysates with Na_3_VO_4_. *, p ≤ 0.05; **, p ≤ 0.01; ***, p ≤ 0.001.

**Supplemental Figure S3.1:**
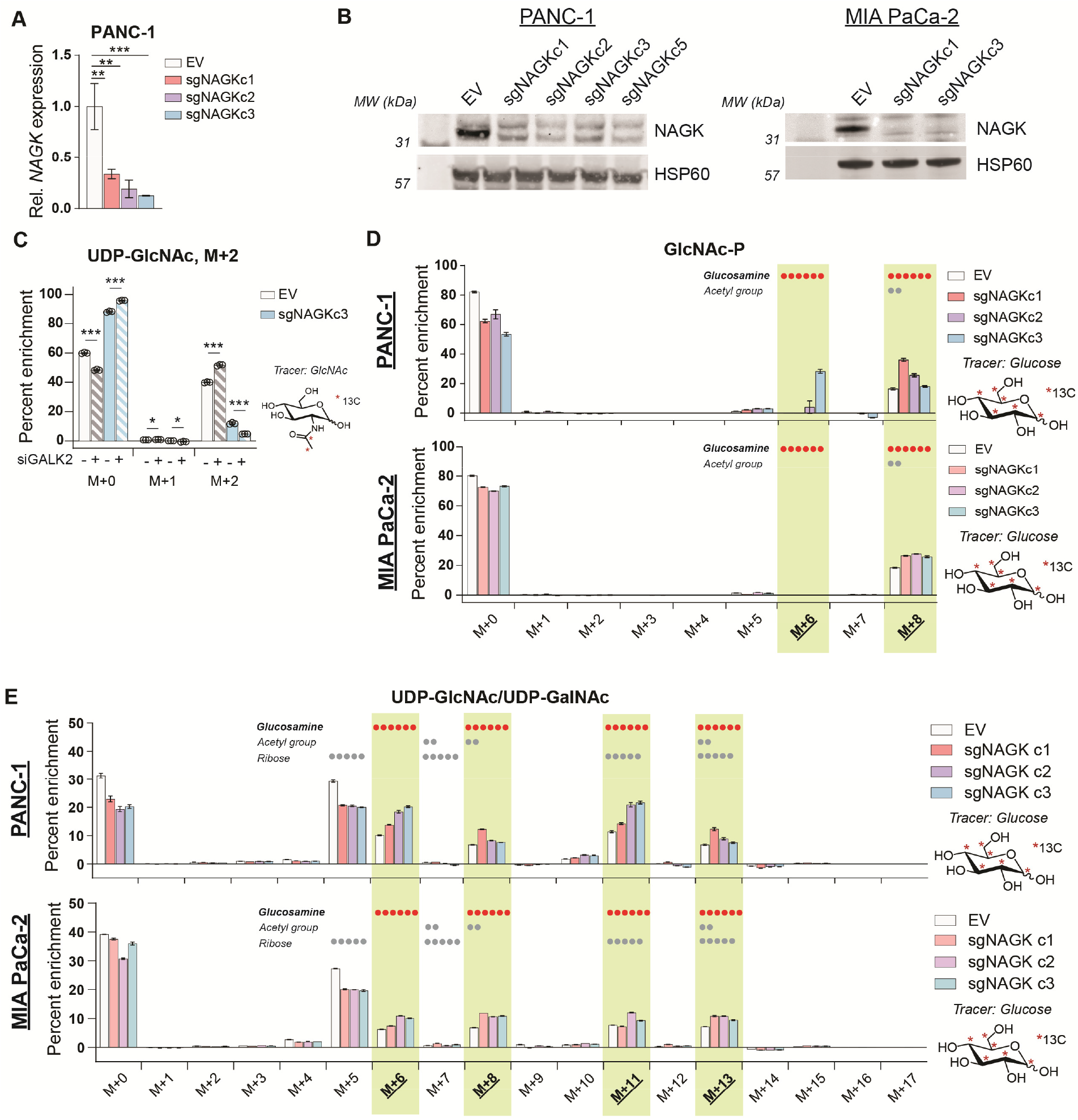
NAGK knockout cells show increased enrichment of 13C glucose into hexosamine intermediates. A) Gene expression of NAGK in PANC-1 NAGK CRISPR cells. Mean +/- SEM of three biological replicates is represented. Statistical significance was determined by one-way ANOVA. **B)** Western blot for NAGK and HSP6O in PANC-1 (left) and MIA PaCa-2 (right) NAGK CRISPR knockout cells. **C)** Measurement of incorporation of 13C GlcNAc labeled on the acetyl group into UDP-GlcNAc in PANC-1 NAGK knockout cells. Cells were incubated with 10 mM 13C GlcNAc for 3 hours. Mean +/- SEM of three biological replicates is represented. Statistical significance was determined by unpaired t-test comparing siCon and siGALK2 for each clone. **D)** Measurement of incorporation of 13C glucose into GlcNAc-P in NAGK knockout cells. Mean +/- SEM of three biological replicates is represented. **E)** Measurement of incorporation of 13C glucose into UDP-GlcNAc in NAGK knockout cells. Mean +/- SEM of three biological replicates is represented. For(D-E), components of molecule labeled in each isotopologue are indicated, with dots representing the number of carbons; highlighted panels identify glucosamine ring labeling. *, p ≤ 0.05; **, p ≤ 0.01; ***, p ≤ 0.01.

**Supplemental Figure S3.2:**
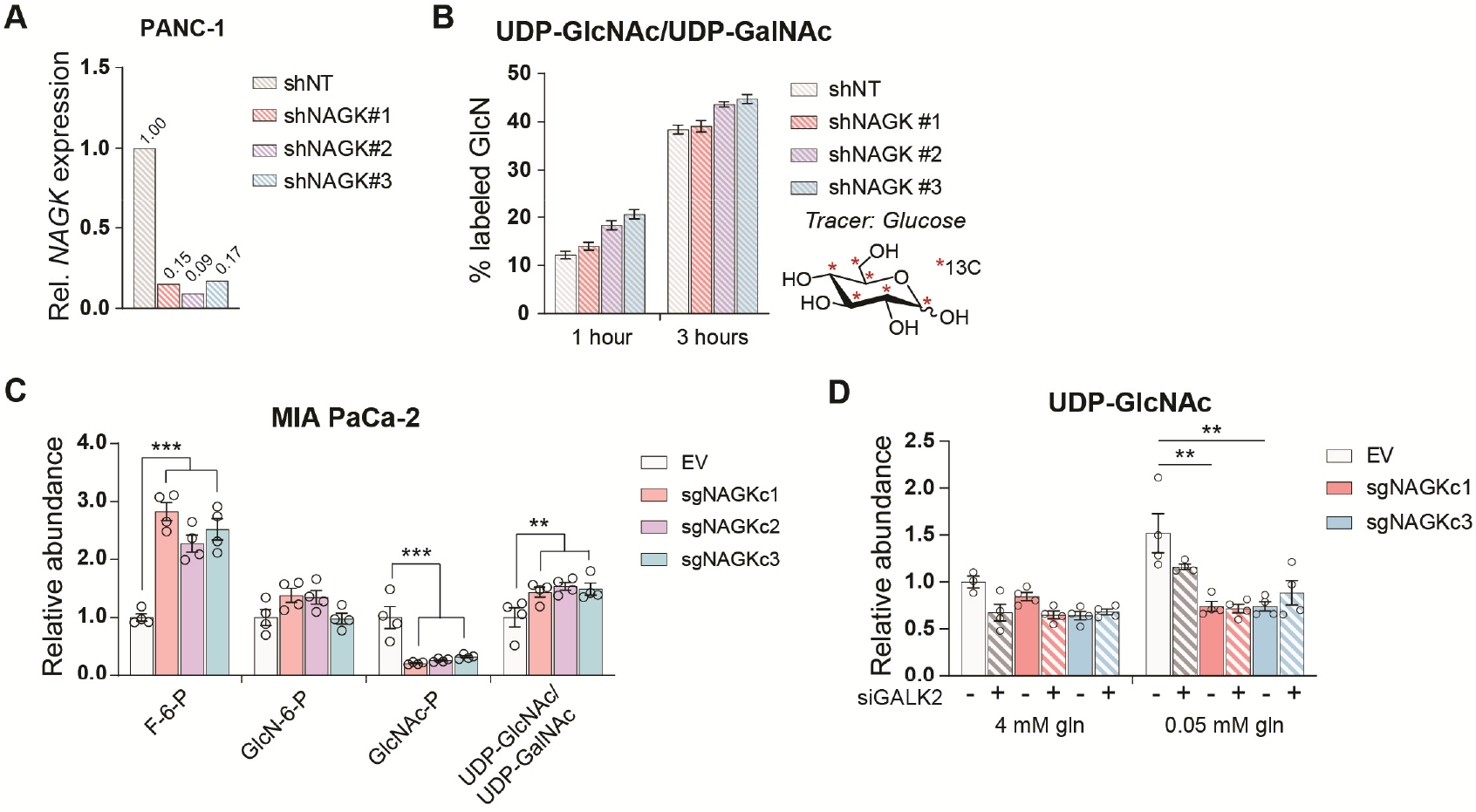
HBP metabolites in NAGK KO cells. **A)** Gene expression of NAGK in PANC-1 cells transfected with shRNA against NAGK. One biological sample is represented. **B)** Incorporation of 13C glucose into the glucosamine ring of UDP-GlcNAc in PANC-1 cells transfected with the indicated shRNAs targeting NAGK. Mean +/- SEM of three biological replicates is represented. **C)** Measurement of HBP metabolites in MIA PaCa-2 NAGK knockout cells cultured in 4 mM glutamine. Mean +/- SEM of four biological replicates is represented. Statistical significance was calculated by unpaired t-test comparing the mean of the three CRISPR clones and the empty vector control. **D)** Measurement of UDP-GlcNAc in PANC-1 NAGK knockout cells. Mean +/- SEM of four biological replicates is represented. Statistical significance was calculated by one-way ANOVA comparing each knockout to the EV control. *, p ≤ 0.05; **, p ≤ 0.01; ***, p≤ 0.001.

**Supplemental Figure 4:**
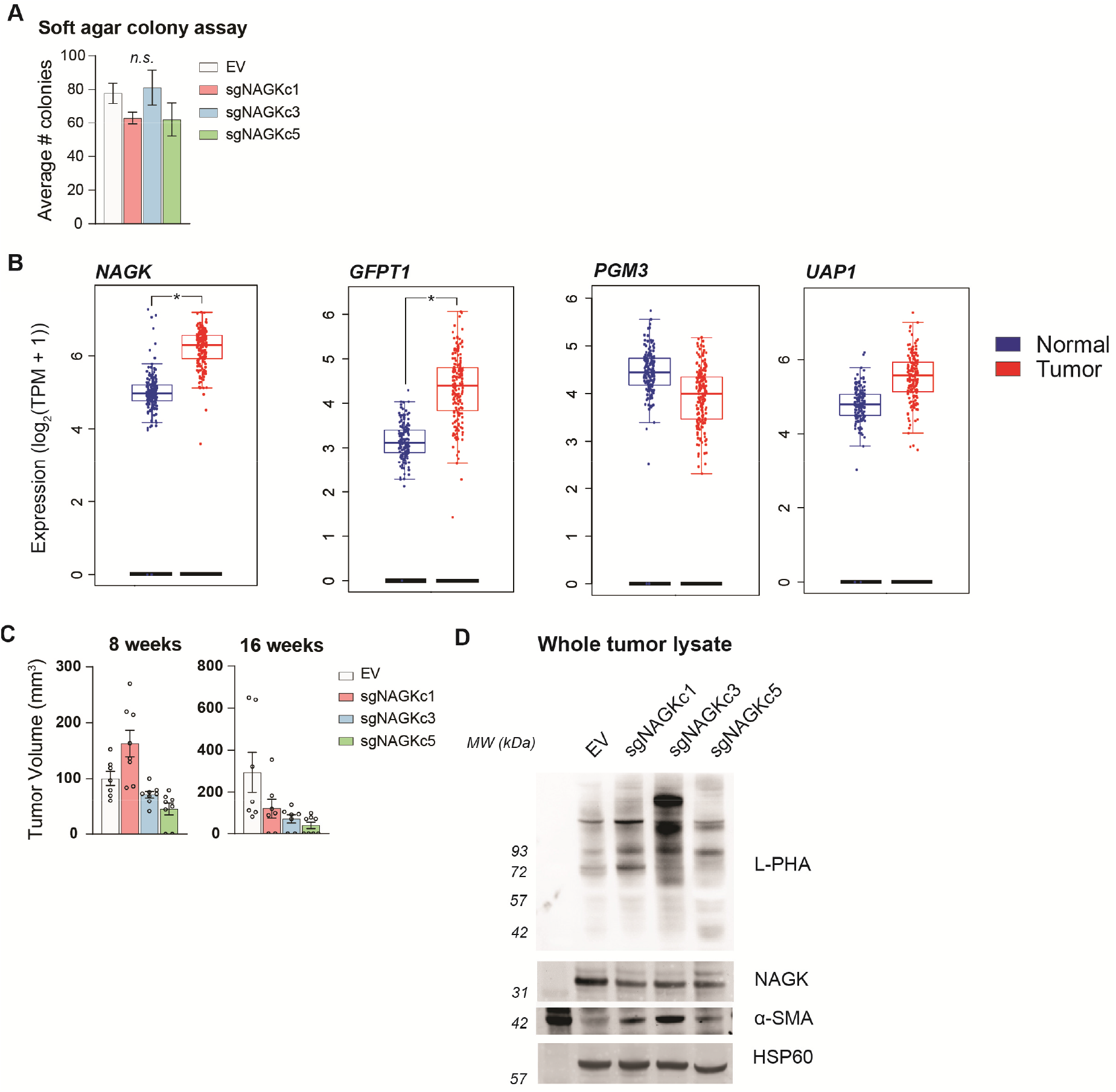
NAGK knockout tumors show increased LPHA binding. **A)** Soft agar colony formation in NAGK knockout and control cells. Mean +/- SEM of three biological replicates is represented. **B)** Gene expression data plotted using GEPIA2 comparing TCGA pancreatic cancer samples with TCGA normal pancreas samples and GTEx data. Differential analysis between tumor and normal tissues was analyzed by one-way ANOVA. *, p < 0.05. **C)** Tumor volume of NAGK knockout tumors at earlier time points in the experiment. Initial growth of NAGK knockout cells in vivo was comparable to control cells, but over time NAGK knockout tumors either stopped growing or regressed. **D)** Western blots for NAGK and ¤-SMA, and blot probed for LPHA binding, in whole tumor lysate samples. Blot for HSP6O as loading control.

## References

Akella, N.M., Ciraku, L., and Reginato, M.J. (2019). Fueling the fire: Emerging role of the hexosamine biosynthetic pathway in cancer. BMC Biol. 17, 1–14.

Caldwell, S.A., Jackson, S.R., Shahriari, K.S., Lynch, T.P., Sethi, G., Walker, S., Vosseller, K., and Reginato, M.J. (2010). Nutrient sensor O-GlcNAc transferase regulates breast cancer tumorigenesis through targeting of the oncogenic transcription factor FoxM1. Oncogene 29, 2831–2842.

Chen, R., Lai, L.A., Sullivan, Y., Wong, M., Wang, L., Riddell, J., Jung, L., Pillarisetty, V.G., Brentnall, T.A., and Pan, S. (2017). Disrupting glutamine metabolic pathways to sensitize gemcitabine-resistant pancreatic cancer. Sci. Rep. 7, 1–14.

Commisso, C., Davidson, S.M., Soydaner-Azeloglu, R.G., Parker, S.J., Kamphorst, J.J., Hackett, S., Grabocka, E., Nofal, M., Drebin, J.A., Thompson, C.B., et al. (2013). Macropinocytosis of protein is an amino acid supply route in Ras-transformed cells. Nature 497, 633–637.

Cutsem, E. Van, Tempero, M.A., Sigal, D., Oh, D., Fazio, N., Macarulla, T., Hitre, E., Hammel, P., Hendifar, A., Bates, S., et al. (2020). Randomized Phase III Trial of Pegvorhyaluronidase Alfa With Nab-Paclitaxel Plus Gemcitabine for Patients With Hyaluronan-High Metastatic Pancreatic Adenocarcinoma abstract. J. Clin. Oncol.

Denzel, M.S., and Antebi, A. (2015). Hexosamine pathway and (ER) protein quality control. Curr. Opin. Cell Biol. 33, 14–18.

Dickinson, M.E., Flenniken, A.M., Ji, X., Teboul, L., Wong, M.D., White, J.K., Meehan, T.F., Weninger, W.J., Westerberg, H., Adissu, H., et al. (2016). High-throughput discovery of novel developmental phenotypes. Nature 537, 508–514.

Doench, J.G., Fusi, N., Sullender, M., Hegde, M., Vaimberg, E.W., Donovan, K.F., Smith, I., Tothova, Z., Wilen, C., Orchard, R., et al. (2016). Optimized sgRNA design to maximize activity and minimize off-target effects of CRISPR-Cas9. Nat. Biotechnol. 34, 184–191.

Edgar, R., Domrachev, M., and Lash, A.E. (2002). Gene Expression Omnibus: NCBI gene expression and hybridization array data repository. Nucleic Acids Res. 30, 207–210.

Engle, D.D., Tiriac, H., Rivera, K.D., Pommier, A., Oni, T.E., Alagesan, B., Lee, E.J., Yao, M.A., Lucito, M.S., Spielman, B., et al. (2019). The Glycan CA19-9 Promotes Pancreatitis and Pancreatic Cancer in Mice. 364, 1156–1162.

Ferrer, C.M., Lu, T.Y., Bacigalupa, Z.A., Katsetos, C.D., Sinclair, D.A., and Reginato, M.J. (2017). O-GlcNAcylation regulates breast cancer metastasis via SIRT1 modulation of FOXM1 pathway. Oncogene 36, 559–569.

Granovsky, M., Fata, J., Pawling, J., Muller, W.J., Khokha, R., and Dennis, J.W. (2000). Suppression of tumor growth and metastasis in Mgat5-deficient mice. Nat. Med. 6, 306–312.

Gu, Y., Mi, W., Ge, Y., Liu, H., Fan, Q., Han, C., Yang, J., Han, F., Lu, X., and Yu, W. (2010). GlcNAcylation plays an essential role in breast cancer metastasis. Cancer Res. 70, 6344–6351.

Guillaumond, F., Leca, J., Olivares, O., Lavaut, M.-N.N., Vidal, N., Berthezène, P., Dusetti, N.J., Loncle, C., Calvo, E., Turrini, O., et al. (2013). Strengthened glycolysis under hypoxia supports tumor symbiosis and hexosamine biosynthesis in pancreatic adenocarcinoma. Proc. Natl. Acad. Sci. U. S. A. 110, 3919–3924.

Guo, H., Zhang, B., Nairn, A. V., Nagy, T., Moremen, K.W., Buckhaults, P., and Pierce, M. (2017). O-Linked N-Acetylglucosamine (O-GlcNAc) Expression Levels Epigenetically Regulate Colon Cancer Tumorigenesis by Affecting the Cancer Stem Cell Compartment via Modulating Expression of Transcriptional Factor MYBL1. J. Biol. Chem. 292, 4123–4137.

Guo, J.Y., Chen, H.Y., Mathew, R., Fan, J., Strohecker, A.M., Karsli-Uzunbas, G., Kamphorst, J.J., Chen, G., Lemons, J.M.S., Karantza, V., et al. (2011). Activated Ras requires autophagy to maintain oxidative metabolism and tumorigenesis. Genes Dev. 25, 460–470.

Guo, L., Worth, A.J., Mesaros, C., Snyder, N.W., Jerry, D., and Blair, I.A. (2016). Diisopropylethylamine/hexafluoroisopropanol-mediated ion-pairing UHPLC-MS for phosphate and carboxylate metabolite analysis: utility for studying cellular metabolism. Rapid Commun Mass Spectrom. 30, 1835–1845.

Halbrook, C.J., and Lyssiotis, C.A. (2017). Employing Metabolism to Improve the Diagnosis and Treatment of Pancreatic Cancer. Cancer Cell 31, 5–19.

Housley, M.P., Rodgers, J.T., Udeshi, N.D., Kelly, T.J., Shabanowitz, J., Hunt, D.F., Puigserver, P., and Hart, G.W. (2008). O-GlcNAc regulates FoxO activation in response to glucose. J. Biol. Chem. 283, 16283–16292.

Jacobetz, M.A., Chan, D.S., Neesse, A., Bapiro, T.E., Cook, N., Frese, K.K., Feig, C., Nakagawa, T., Caldwell, M.E., Zecchini, H.I., et al. (2013). Hyaluronan impairs vascular function and drug delivery in a mouse model of pancreatic cancer. Gut 62, 112–120.

Kamphorst, J.J., Nofal, M., Commisso, C., Hackett, S.R., Lu, W., Grabocka, E., Vander Heiden, M.G., Miller, G., Drebin, J.A., Bar-Sagi, D., et al. (2015). Human Pancreatic Cancer Tumors Are Nutrient Poor and Tumor Cells Actively Scavenge Extracellular Protein. Cancer Res. 75, 544–553.

Kim, P.K., Halbrook, C.J., Kerk, S.A., Wisner, S., Kremer, D., Sajjakulnukit, P., Hou, S.W., Thurston, G., Anand, A., Yan, L., et al. (2020). Hyaluronic Acid Fuels Pancreatic Cancer Growth. BioRxiv.

Lau, K.S., Partridge, E.A., Grigorian, A., Silvescu, C.I., Reinhold, V.N., Demetriou, M., and Dennis, J.W. (2007). Complex N-Glycan Number and Degree of Branching Cooperate to Regulate Cell Proliferation and Differentiation. Cell 129, 123–134.

Li, D., Li, Y., Wu, X., Li, Q., Yu, J., Gen, J., and Zhang, X.-L. (2008). Knockdown of Mgat5 Inhibits Breast Cancer Cell Growth with Activation of CD4 + T Cells and Macrophages. J. Immunol. 180, 3158–3165.

Lynch, T.P., Ferrer, C.M., Jackson, S.R.E., Shahriari, K.S., Vosseller, K., and Reginato, M.J. (2012). Critical role of O-linked β-N-acetylglucosamine transferase in prostate cancer invasion, angiogenesis, and metastasis. J. Biol. Chem. 287, 11070–11081.

Lyssiotis, C.A., and Kimmelman, A.C. (2017). Metabolic Interactions in the Tumor Microenvironment. Trends Cell Biol. 27, 863–875.

Mahlbacher, V., Sewing, A., Elsässer, H.P., and Kern, H.F. (1992). Hyaluronan is a secretory product of human pancreatic adenocarcinoma cells. Eur. J. Cell Biol. 58, 28–34.

Mereiter, S., Balmaña, M., Campos, D., Gomes, J., and Reis, C.A. (2019). Glycosylation in the Era of Cancer-Targeted Therapy: Where Are We Heading? Cancer Cell 36, 6–16.

Moloughney, J.G.G., Kim, P.K.K., Vega-cotto, N.M.M., Wu, C.-C., Zhang, S., Adlam, M., Lynch, T., Chou, P.-C., Rabinowitz, J.D.D., Werlen, G., et al. (2016). mTORC2 Responds to Glutamine Catabolite Levels to Modulate the Hexosamine Biosynthesis Enzyme GFAT1. Mol. Cell 63, 811–826.

Moseley, H.N.B., Lane, A.N., Belshoff, A.C., Higashi, R.M., and Fan, T.W.M. (2011). A novel deconvolution method for modeling UDP-N-acetyl-D-glucosamine biosynthetic pathways based on 13C mass isotopologue profiles under non-steady-state conditions. BMC Biol. 9.

Munkley, J. (2019). The glycosylation landscape of pancreatic cancer (Review). Oncol. Lett. 17, 2569–2575.

Özdemir, B.C., Pentcheva-Hoang, T., Carstens, J.L., Zheng, X., Wu, C.C., Simpson, T.R., Laklai, H., Sugimoto, H., Kahlert, C., Novitskiy, S. V., et al. (2014). Depletion of carcinoma-associated fibroblasts and fibrosis induces immunosuppression and accelerates pancreas cancer with reduced survival. Cancer Cell 25, 719–734.

Park, S.Y., Kim, H.S., Kim, N.H., Ji, S., Cha, S.Y., Kang, J.G., Ota, I., Shimada, K., Konishi, N., Nam, H.W., et al. (2010). Snail1 is stabilized by O-GlcNAc modification in hyperglycaemic condition. EMBO J. 29, 3787–3796.

Pei, H., Li, L., Fridley, B.L., Jenkins, G.D., Kalari, K.R., Lingle, W., Petersen, G., Lou, Z., and Wang, L. (2009). FKBP51 Affects Cancer Cell Response to Chemotherapy by Negatively Regulating Akt. Cancer Cell 16, 259–266.

Provenzano, P.P., Cuevas, C., Chang, A.E., Goel, V.K., Von Hoff, D.D., and Hingorani, S.R. (2012). Enzymatic Targeting of the Stroma Ablates Physical Barriers to Treatment of Pancreatic Ductal Adenocarcinoma. Cancer Cell 21, 418–429.

Rahib, L., Smith, B.D., Aizenberg, R., Rosenzweig, A.B., Fleshman, J.M., and Matrisian, L.M. (2014). Projecting cancer incidence and deaths to 2030: The unexpected burden of thyroid, liver, and pancreas cancers in the united states. Cancer Res. 74, 2913–2921.

Ricciardiello, F., Votta, G., Palorini, R., Raccagni, I., Brunelli, L., Paiotta, A., Tinelli, F., D’Orazio, G., Valtorta, S., De Gioia, L., et al. (2018). Inhibition of the Hexosamine Biosynthetic Pathway by targeting PGM3 causes breast cancer growth arrest and apoptosis. Cell Death Dis. 9.

Ricciardiello, F., Gang, Y., Palorini, R., Li, Q., Giampà, M., Zhao, F., You, L., La Ferla, B., De Vitto, H., Guan, W., et al. (2020). Hexosamine pathway inhibition overcomes pancreatic cancer resistance to gemcitabine through unfolded protein response and EGFR-Akt pathway modulation. Oncogene 4103–4117.

Ryczko, M.C., Pawling, J., Chen, R., Abdel Rahman, A.M., Yau, K., Copeland, J.K., Zhang, C., Surendra, A., Guttman, D.S., Figeys, D., et al. (2016). Metabolic Reprogramming by Hexosamine Biosynthetic and Golgi N-Glycan Branching Pathways. Sci. Rep. 6, 23043.

Sanjana, N.E., Shalem, O., and Zhang, F. (2014). Sanjana, Shalem et Zhang. Nat. Med. 11, 783–784.

Schneider, C.A., Rasband, W.S., and Eliceiri, K.W. (2012). NIH Image to ImageJ: 25 years of image analysis. Nat. Methods 9, 671–675.

Sharma, N.S., Saluja, A., Banerjee, S., Sharma, N.S., Gupta, V.K., Garrido, V.T., Hadad, R., Durden, B.C., Kesh, K., Giri, B., et al. (2020). Targeting tumor-intrinsic hexosamine biosynthesis sensitizes pancreatic cancer to anti-PD1 therapy Graphical abstract Find the latest version: Targeting tumor-intrinsic hexosamine biosynthesis sensitizes pancreatic cancer to anti-PD1 therapy. 130, 451–465.

Steenackers, A., Olivier-Van Stichelen, S., Baldini, S.F., Dehennaut, V., Toillon, R.A., Le Bourhis, X., El Yazidi-Belkoura, I., and Lefebvre, T. (2016). Silencing the nucleocytoplasmic O-GlcNAc transferase reduces proliferation, adhesion, and migration of cancer and fetal human colon cell lines. Front. Endocrinol. (Lausanne). 7, 1–11.

Swamy, M., Pathak, S., Grzes, K.M., Damerow, S., Sinclair, L. V, van Aalten, D.M.F., and Cantrell, D.A. (2016). Glucose and glutamine fuel protein O-GlcNAcylation to control T cell selfrenewal and malignancy. Nat. Immunol. 17, 1–11.

Tang, Z., Kang, B., Li, C., Chen, T., and Zhang, Z. (2019). GEPIA2: an enhanced web server for large-scale expression profiling and interactive analysis. Nucleic Acids Res. 47, W556–W560.

Taylor, R.P., Parker, G.J., Hazel, M.W., Soesanto, Y., Fuller, W., Yazzie, M.J., and McClain, D.A. (2008). Glucose deprivation stimulates O-GlcNAc modification of proteins through upregulation of O-linked N-acetylglucosaminyltransferase. J. Biol. Chem. 283, 6050–6057.

Trefely, S., Ashwell, P., and Snyder, N.W. (2016). FluxFix: Automatic isotopologue normalization for metabolic tracer analysis. BMC Bioinformatics 17, 1–8.

Wellen, K.E., Lu, C., Mancuso, A., Lemons, J.M.S.S., Ryczko, M., Dennis, J.W., Rabinowitz, J.D., Coller, H.A., and Thompson, C.B. (2010). The hexosamine biosynthetic pathway couples growth factor-induced glutamine uptake to glucose metabolism. Genes Dev. 24, 2784–2799.

Wyant, G.A., Abu-Remaileh, M., Frenkel, E.M., Laqtom, N.N., Dharamdasani, V., Lewis, C.A., Chan, S.H., Heinze, I., Ori, A., and Sabatini, D.M. (2018). NUFIP1 is a ribosome receptor for starvation-induced ribophagy. Science (80-.). 360, 751–758.

Ying, H., Kimmelman, A.C., Lyssiotis, C.A., Hua, S., Chu, G.C., Fletcher-Sananikone, E., Locasale, J.W., Son, J., Zhang, H., Coloff, J.L., et al. (2012). Oncogenic kras maintains pancreatic tumors through regulation of anabolic glucose metabolism. Cell 149, 656–670.

Zhou, X., Chen, H., Wang, Q., Zhang, L., and Zhao, J. (2011). Knockdown of Mgat5 inhibits CD133+ human pulmonary adenocarcinoma cell growth in vitro and in vivo. Clin. Invest. Med. 34, E155–62.

